# Bathymetric Resolution-Dependent Biases in Antarctic Benthic Biodiversity Models: Hotspots Hold, Counts Shift

**DOI:** 10.64898/2026.06.23.734136

**Authors:** Shaylyn R. Potter, Jan Jansen, Nicole Hill, Vanessa Lucieer

## Abstract

Antarctic benthic organisms are highly diverse and play a critical role in the Southern Ocean ecosystem. Despite decades of sampling, vast areas of the Antarctic continental shelf remain biologically unsurveyed due to logistical and financial constraints, limiting baseline knowledge essential for effective conservation planning. Species distribution models (SDMs) allow biodiversity to be inferred in the absence of biological data by linking benthic community patterns to environmental predictors. However, the resolution of the environmental predictors, particularly bathymetry, varies significantly between regions, casting doubt about how reliably SDMs can be used to predict into regions where only coarse-resolution data are available. Here, we show that SDMs trained on high-resolution data underestimate Antarctic benthic morphospecies richness by up to 18% when applied to aggregated coarse-resolution environmental data (and up to 50% when using satellite-derived ETOPO bathymetry). Using six systematically degraded versions of high-resolution multibeam bathymetry and annotated seafloor imagery across three Antarctic regions, we evaluate SDM performance both with and without additional environmental variables. High-resolution bathymetry captures terrain complexity most effectively, but we find that the spatial distribution of richness hotspots and the median richness per cell remain consistent, provided models are applied at the same resolution at which they were trained. Our results suggest that while high-resolution bathymetry may enhance local predictions, coarse-resolution data may be more robust for regional-scale predictions, such as those used for Antarctic shelf-wide spatial planning.

## Introduction

The Southern Ocean hosts a rich and highly endemic benthic fauna, which plays a vital role in maintaining the function and resilience of Antarctic marine ecosystems (De Broyer et al. 2011; Griffiths et al. 2009; Barnes and Tarling 2017). Vast areas of the Antarctic seafloor remain biologically under-surveyed due to logistical and financial constraints (De Broyer and Koubbi 2014), especially in remote regions such as the Eastern Ross Sea, Western Weddell Sea, East Antarctica, and depths greater than 500 m (Dorschel et al. 2022). As climate change and human pressures on the Southern Ocean intensify (Constable et al. 2014; Griffiths et al. 2024), benthic organisms will be directly affected (Amsler et al. 2023; Barnes and Peck 2008). Understanding species distributions and identifying biodiversity hotspots has therefore become a priority for Antarctic marine spatial planning and management (Brasier et al. 2021).

The long-recognised affinity of benthic fauna to specific environmental conditions and depths (Džeroski and Drumm 2003; Gutt and Starmans 1998; Post et al. 2017; Almond et al. 2021), demonstrates that abiotic information can be used to infer biotic patterns when biological data are scarce. Terrain variables (e.g. slope, rugosity) derived from bathymetric data and other environmental data (e.g. temperature, currents, food availability) can be linked to biological information from georeferenced imagery through statistical models to identify biodiversity hotspots and predict species distributions (Dolan and Lucieer 2014; Lucieer et al. 2013). In this way, predictive models, particularly species distribution models (SDMs), provide a valuable tool for estimating biodiversity in unsurveyed regions and support large-scale management (Gros et al. 2022; Peterson and Herkül 2019). These models have been widely applied in marine systems, including coral reefs (Harris et al. 2013), temperate rocky reefs (Hill et al. 2014), the deep sea (Rengstorf et al. 2014; Winship et al. 2020), continental shelves (Kostylev et al. 2001), and across the Antarctic seafloor (Jansen et al. 2018; Hogg et al. 2018; Smith et al. 2015).

The reliability of SDMs, however, depends heavily on the resolution and quality of spatial predictors, such as bathymetry (Misiuk et al. 2021; Lecours et al. 2015). While high-resolution bathymetric data are optimal for detecting habitat-forming features (Post et al. 2020; Ross et al. 2015) as well as habitat complexity (Leite Jardim et al. 2025), high-resolution, or small spatial scale, does not always yield better model performance. Some studies suggest intermediate or coarser environmental data resolutions may be more effective to capture regional-scale processes (Monk et al. 2011; Porskamp et al. 2018; Núñez-Riboni et al. 2021). Bathymetric grids across the Southern Ocean are compiled from heterogeneous datasets that vary in quality, scale, and acquisition methods. According to the International Bathymetric Chart of the Southern Ocean (IBCSO: Dorschel et al. 2022), only 22.32% of the region is covered by multibeam echosounder (MBES) data, while 1.47% comes from single-beam or other soundings, leaving more than 76% derived from low-resolution satellite altimetry. The transition from IBCSO version 1 (Arndt et al. 2013) to version 2 (Dorschel et al. 2022) highlighted large discrepancies in seafloor representation across different resolutions, especially in steep-slope regions where MBES data replaced satellite data. These discrepancies matter because, with the Antarctic shelf ecosystems exhibiting both fine-scale habitat heterogeneity (Post et al. 2017; Ziegler et al. 2017) and regional-scale influences (Dawson et al. 2023), it remains unclear what bathymetric resolution is needed to both capture key seafloor features and produce robust shelf-wide models.

Despite the recognised importance of bathymetry in predictive modelling, there is limited understanding of how mismatched resolutions between training and prediction datasets bias biodiversity estimates. In large compilations such as IBCSO, fine-scale MBES data are included but aggregated to the resolution of the overall product, and imagery data are often concentrated in areas where high-resolution bathymetry is available. This means models are frequently developed on high-resolution bathymetry but then applied to predict biodiversity patterns across broader regions represented only by coarse-resolution data. The key question is whether such resolution mismatches lead to systematic under- or overestimation of biodiversity patterns.

To address this, we test how varying bathymetric resolution affects predictive models of habitat suitability and biodiversity across three Antarctic shelf regions. Using systematically degraded versions of high-resolution bathymetry and coinciding annotated imagery (Jansen et al. 2026), we evaluate changes in terrain attributes, model performance (both with and without the inclusion of additional environmental variables), predictive maps, and test how predictive models trained on high-resolution bathymetry perform when applied to coarser-resolution data. This research provides insight into the trade-offs between data availability and predictive performance, informing future SDM applications in under-surveyed regions of the Southern Ocean.

## Methods

### Study Areas

The Antarctic continental shelf, shaped by its glacial history, is deeper (average 450 m, exceeding 1,000 m in some areas) and wider (∼125 km) than global averages (Clarke and Johnston 2003; Smith et al. 2006), with sediments composed primarily of diatomaceous muds and glacial deposits (Clarke 1996; Griffiths 2010). This study focused on three ecologically distinct regions along the Antarctic continental shelf: the Sabrina Coast (SC), the Ross Sea (RS), and the Antarctic Peninsula (AP) (Figure 1).

**Figure 1.**
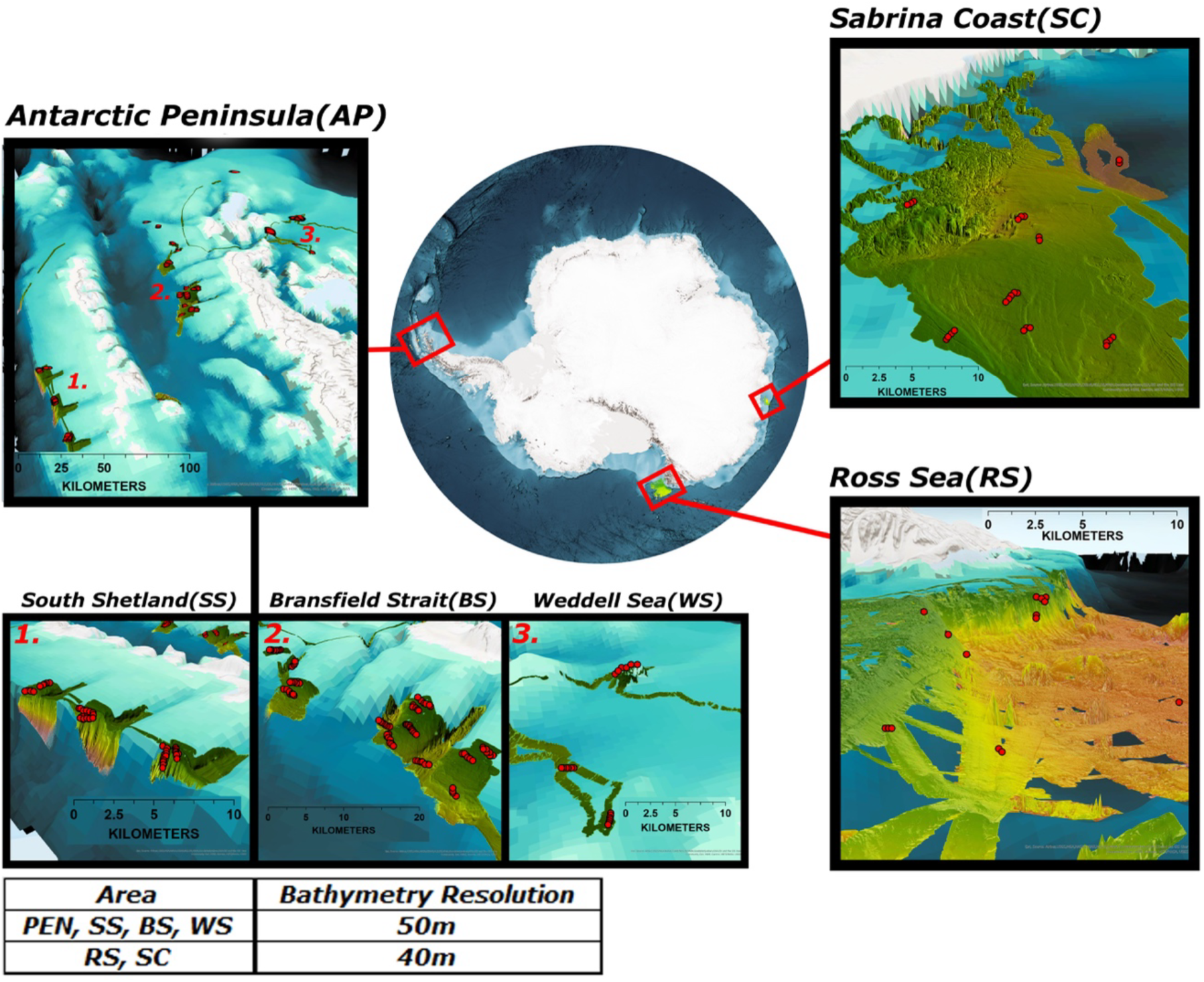
Overview map of study area. Subset images show the multibeam bathymetric data used with marked coordinates (red dots) of seafloor image locations. Study areas are the Sabrina Coast (SC), Ross Sea (RS), and the Antarctic Peninsula (AP). The Peninsula study area was split into three different regions 1. South Shetland (SS) 2. Bransfield Strait (BS) 3. Weddell Sea (WS) for some analyses.

The SC region, located between the Moscow University Ice Shelf and the Totten Glacier, features irregular over-deepened bathymetry sculpted by glacial erosion (Montelli et al. 2020), with prominent features including bedrock with complex channels that deepen as it progresses to the shelf break and the Aurora Channel.

The Ross Sea dataset (RS) was collected in the northwest Ross Sea, just off the north tip of the Adare Peninsula. Surrounding this cape, the continental shelf greatly narrows and quickly drops over the continental slope and into the Adare Basin. Prominent seafloor features include the continental shelf, JOIDES Basin, Adare Basin, Hallett Ridge, and Adare Trough (Selvans et al. 2014). This study site falls partially within the boundaries of the Ross Sea Marine Protected Area (MPA; Conservation Measure 91-05, 2016).

The AP region features strong environmental gradients (Dorschel et al. 2014), complex seafloor topography (Dorschel et al. 2022; Jerosch et al. 2016), and high biodiversity (Clarke and Johnston 2003; Griffiths 2010), influenced by the mixing of Circumpolar Deep Water and Weddell Sea Water, and variable sea ice and productivity (Clarke et al. 2009; Zhou et al. 2006). It includes three ecoregions (Gutt et al. 2016): (1) Drake Passage, with shallow shelves and canyon systems; (2) Bransfield Strait, with flat shelves and coastal canyons; and (3) the Western Weddell Sea, with smoother slopes and higher sea ice cover.

### Software and Packages

*RStudio* (2023) was used for visualising raster plots, all raster and terrain attribute calculations, statistical analyses, and creation of predictive maps. Packages loaded and used included: *terra* v1.8-67 (Hijmans 2023)*, MASS* version 7.3-58.1 (Venables and Ripley 2002), *PerformanceAnalytics* version 2.0 (Peterson and Carl 2020), *dplyr* version 1.0.10 (Wickham et al. 2022)*, ggplot2* version 3.4.0 (Wickham 2016), and *lme4* version 1.1-31 (Bates et al. 2015).

### Environmental data

#### Bathymetry

High-resolution bathymetric data from multibeam echosounder (MBES) surveys were identified from the International Bathymetric Chart of the Southern Ocean (IBCSO; Dorschel et al. 2022; https://ibcso.org) for areas that aligned with available biological imagery. We sourced original bathymetric surveys from the data custodians, gridded at native resolutions of 40 or 50 m, and systematically degraded them to 100 m, 500 m, 1000 m, 2000 m, and 4000 m to simulate commonly encountered datasets in predictive modelling. Details of each survey, including institution and collection year, are listed in Table 1. As a comparison, we included ETOPO satellite-derived bathymetry from the 1 Arc-Minute Global Relief Model (Amante and Eakins 2009) that has a working resolution of approximately 500–600 m when projected to Southern Ocean coordinates (EPSG:9354). Since ETOPO predates the inclusion of modern MBES data (Tozer et al. 2019), it served as a representation of purely satellite-derived bathymetry to assess how well our degraded rasters simulate coarse-resolution global data.

**Table 1:** Publicly available environmental and biological data used in the study with associated years, institutions, and survey IDs.

| Type of data | Data collection | Year collected | Institution | Region | Survey ID |
| --- | --- | --- | --- | --- | --- |
| Environmental<br>(bathymetric) | MBES | 2014 | NCEI | Sabrina Coast | NBP1402<br>IBCSO file:<br>NBP1402.xyz |
| Environmental<br>(bathymetric) | MBES | 2006/2007 | NCEI | Ross Sea West | NBP0701<br>IBCSO file:<br>NBP0701.xyz |
| Environmental<br>(bathymetric) | MBES | 2013 | AWI | Peninsula | PS81<br>IBCSO file:<br>Ant29_3.xyz |
| Environmental<br>(bathymetric) | Satellite<br>interpolated | Prior to 2009 | ETOPO1<br>Amante &<br>Eakins, 2009 | Circumpolar | N/A |
| Environmental<br>(NPP) | Satellite<br>CAFE model | 2002-2014 | Silsbe et al.,<br>2016 | Circumpolar | N/A |
| Environmental<br>(seafloor currents) | Numerical<br>model<br>WAOM | Simulated 2007 | Boeira-Dias et<br>al., 2023 | Circumpolar | N/A |
| Environmental<br>(seafloor<br>temperature) | Numerical<br>model<br>WAOM | Simulated 2007 | Boeira-Dias et<br>al., 2023 | Circumpolar | N/A |
| Biological | Yo-Yo camera | 2014 | NCEI | Sabrina Coast | NBP1402 |
| Biological | DTIS | 2008 | NIWA | Ross Sea West | TAN0802 |
| Biological | DTIS | 2019 | NIWA | Ross Sea West | TAN1901 |
| Biological | OFOS | 2013 | AWI | Antarctic Peninsula | PS81 |

**Abbreviations include:** Multibeam echosounder (MBES), National Centers for Environmental Information (NCEI), Alfred Wegener Institute (AWI), International Bathymetric Chart of the Southern Ocean (IBCSO), Whole Antarctic Ocean Model (WAOM), Carbon Absorption and Fluorescence Euphotic-resolving (CAFÉ), Deep Towed Imaging System (DTIS), Ocean Floor Observation System (OFOS), National Institute of Water and Atmospheric Research (NIWA).

#### Bathymetric Data Processing and Deriving Terrain Variables

The SC, RS, and AP bathymetric datasets were imported into R as rasters and aggregated (hereafter, ‘degraded’) to the five coarser resolutions using the *aggregate()* function in the *terra* package (Hijmans 2022). Raster aggregation occasionally resulted in missing cell values (NAs), so we applied a mask to remove cells in higher-resolution rasters not retained at coarser resolutions. Depth and slope were selected as terrain variables for analysis; roughness and terrain ruggedness index (TRI) were excluded due to high collinearity with slope (r > 0.95). All terrain layers were projected to the IBCSO WGS 84 Polar Stereographic CRS (EPSG:9354). To ensure comparability across resolutions, we standardised raster extents and cell counts using the *resample()* function *(method = “near”)*. This enabled consistent cell-wise comparisons and spatial stacking across all resolutions. In the Ross Sea, the native 40 m bathymetry contained sampling artefacts where abrupt depth changes along some transect lines produced false readings of steep slopes (Appendix S1: Figure S1). Distributional changes in slope and depth with resolution degradation were examined using raster plots with density curves (Figure 2; Appendix S1: Figures S2, S3), and correlation analyses (native resolution vs. 4000 m resolution: Appendix S1: Figures S4, S5).

**Figure 2.**
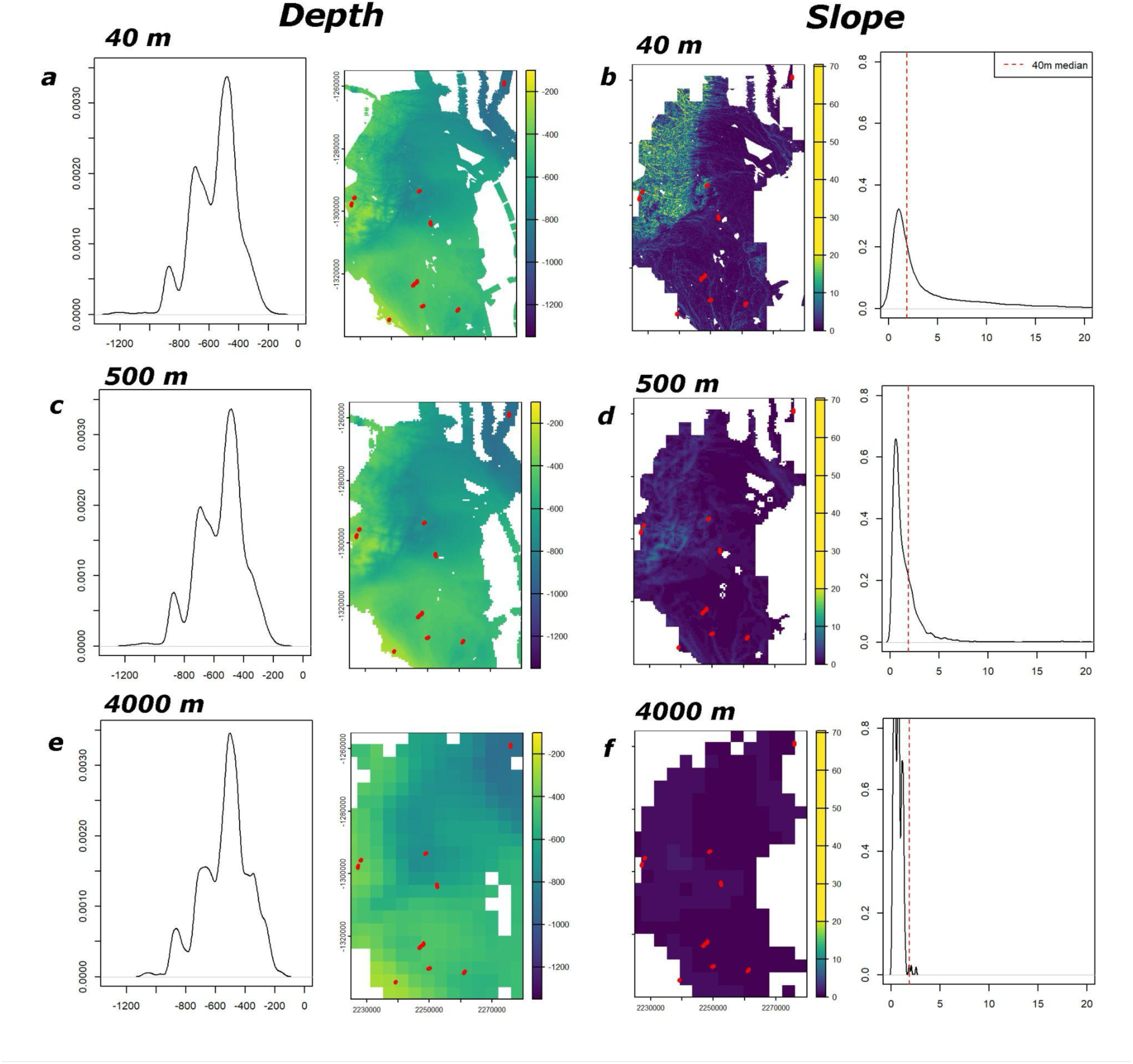
Impact of degrading resolution on depth and slope. Terrain variables depth (**left and centre-left column**) and slope (**centre-right and right column**) of the Sabrina Coast, with density curves (**left and right columns**) comparing the native 40 m raster to aggregated 500 m and 4000 m rasters. Red dots indicate biological sampling locations. Red line on slope density plots (**right column**) represents the median slope value of the native 40 m raster. Depth values change minimally, while slope values flatten significantly, especially at slopes >10°.

#### Additional Environmental Variables

To enhance model performance and test whether environmental predictors could mitigate the effects of bathymetric resolution loss, we incorporated additional variables previously shown to influence benthic species distributions (Jansen et al. 2018). Using environmental predictors alongside terrain variables is a common approach in biodiversity modelling and enables assessment of their relative contributions across resolutions.

The additional variables we included were net primary production (NPP), mean seafloor current speed, and mean seafloor temperature. NPP values were derived from the Carbon, Absorption, and Fluorescence Euphotic-resolving (CAFÉ) model, which uses satellite ocean colour data integrated with ecological and physiological parameters to estimate global productivity (Silsbe et al. 2016). Seafloor currents and temperatures were obtained from a 2 km resolution tide-resolving circumpolar Antarctic Ocean model (Dias et al. 2023). Details can be found in Table 1.

### Biological Data

The biological data used in this study were drawn from a subset of seafloor images curated in the Antarctic Seafloor Annotated Imagery Database (Jansen et al. 2026). This publicly available reference library contains 3,599 annotated images collected from 21 multinational Antarctic research expeditions conducted between 1985 and 2019. All images were acquired using downward-facing sampling platforms to maintain consistency, and only those coinciding spatially with high-resolution multibeam bathymetric surveys were selected for this study (Table 1).

Each image served as a sampling unit and was annotated using a 108-point grid system. Substrate or biota at the centre of each grid point were identified, providing a measure of morpho-group abundance. Annotations were carried out manually with support from CoralNet’s deep-learning-assisted interface (Beijbom et al. 2015) and classified using the CATAMI hierarchical scheme, which balances taxonomic and morphological resolution depending on the level of identification possible (Althaus et al. 2015). All annotations underwent a three-stage quality control process to ensure consistency and accuracy. Further details of image scoring protocols are described in Jansen et al. (2026). The information extracted from the Antarctic Seafloor Annotated Imagery Database included geographic coordinates for each image, survey metadata, the number of taxa recorded in each image (considered here as a metric of species richness and proxy for biodiversity), and total percentage cover of all living organisms. We used percent cover only in supplementary models to confirm that results were not sensitive to the choice of biodiversity metric.

### Analysis

To assess how bathymetric resolution affects predictive modelling of species diversity, we built statistical models for each of the three Antarctic study regions using six bathymetric resolutions (40/50 m, 100 m, 500 m, 1000 m, 2000 m, and 4000 m), plus the satellite-derived ETOPO dataset. Using Generalized Linear Models (GLMs) with a negative binomial distribution, we modelled species richness as the response variable. The terrain attributes depth, its quadratic term (depth²), and log-transformed slope were used as the initial set of predictors.

Models were first run using only bathymetric terrain predictors (depth, slope) to evaluate the independent contribution of bathymetry. We then repeated model fitting with additional environmental predictors: net primary production (NPP), seafloor current speed, and bottom temperature. Due to multicollinearity in the Sabrina Coast (SC) and Ross Sea (RS) datasets (correlation > 0.7 between currents, temperature, and NPP), only NPP was retained alongside the bathymetric predictors for those regions. All environmental variables were included in the Antarctic Peninsula dataset, where no high correlations were detected. For each model, diagnostic plots were used to assess fit, and outputs were evaluated by comparing deviance explained (Figure 3, Appendix S1: Figure S6) and standardised coefficient estimates (Appendix S1: Figure S7). Moving forward, due to stronger model fit in some regions and to keep consistency, predictive maps were generated only from models that included the additional environmental variables. Models were initially fitted separately for the three Antarctic Peninsula ecoregions (Appendix S1: Figures S6, S7), but subsequent analyses used the AP as a single dataset, as pooled models yielded higher deviance explained (DE) values and provided a sufficient and streamlined basis for predictive mapping.

**Figure 3.**
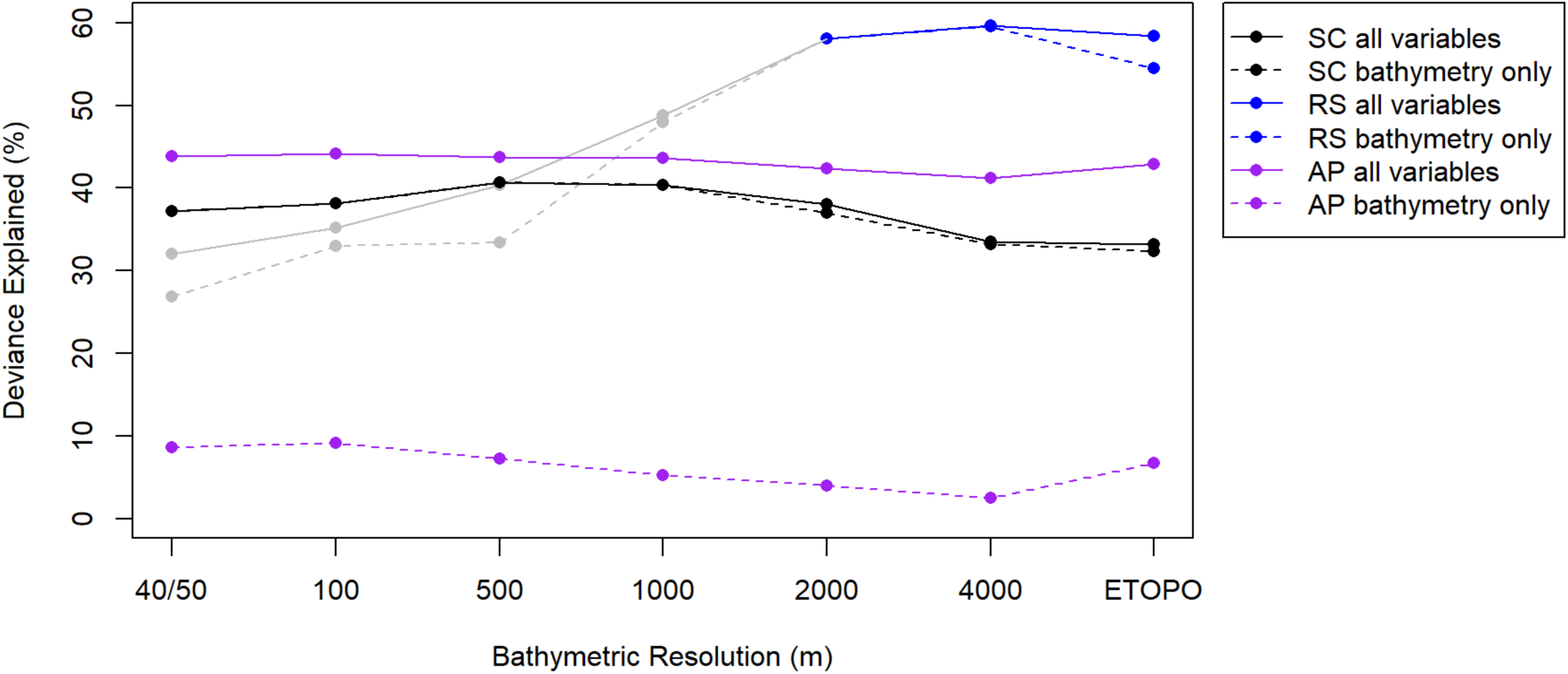
Deviance explained (%) for all resolutions of the datasets. Outputs from the Sabrina Coast (SC), Ross Sea (RS), and Antarctic Peninsula (AP) regions showing deviance explained (%) from Generalized Linear Models using either bathymetric predictors alone or models including additional environmental variables. Resolution changes had little effect on model performance, except in the Ross Sea dataset where sampling artefacts in high-resolution data reduced fit (grey). Key trend is the strong improvement in AP models with added environmental variables.

Data were scaled to help models converge and allow interpretation of coefficients in the relative sense. While we acknowledge that data are clustered, attempts to fit GLMMs, including transect as a random factor, were unsuccessful and therefore the confidence intervals of coefficients, and consequently the significance of any effects, need to be interpreted cautiously (Dormann et al. 2007).

### Predictive Mapping

To visualise and assess how model predictions respond to changes in bathymetric resolution, we generated gridded raster maps of predicted richness for each of the six bathymetry resolutions (40/50 m to 4000 m and ETOPO) in each study region. In addition to fitting and predicting with each model at its own resolution, we explored model transferability by applying the 40 m and 500 m models to environmental predictor layers at each of the other resolutions. This allowed us to isolate how prediction outputs are influenced by resolution differences in the environmental data rather than model coefficients. For each resolution, we calculated the median predicted richness per cell and compared these values across maps to assess trends in over- and underestimation.

To further investigate spatial prediction differences, we calculated richness percentiles (75th, 90th, 95th, and 99th) and compared their spatial distribution across models. Visualisations were created using the *terra* and *ggplot2* packages in *RStudio* (2023). These plots were used to evaluate whether richness hotspots remained consistent in space and to determine how the magnitude of predictions changed with coarser bathymetric inputs. ETOPO-based maps were included to evaluate whether satellite-derived bathymetry yielded similar spatial predictions as degraded multibeam data at comparable resolutions. To confirm that our results were not dependent on the biodiversity metric chosen, we repeated all analyses using percent cover instead of richness.

## Results

### The effect of raster degradation on seafloor terrain attributes

#### Depth

Average raster depth decreases with coarser resolution across all study areas. Maximum depths are shallower and minimum depths deeper as resolution degrades, leading to a narrowing of the overall depth range (Figure 2; Appendix S1: Figures S2, S3, Table S1). Despite this compression, correlation analysis between the native-resolution depth raster (40 m for Sabrina Coast and Ross Sea; 50 m for Antarctic Peninsula) and the coarsest resolution raster (4000 m) shows strong agreement: Sabrina Coast Pearson’s r = 0.94, Ross Sea r = 0.99, Antarctic Peninsula r = 0.95. Full correlation biplots and histograms for each region are provided in the appendix (Appendix S1: Figure S4).

#### Slope

Slope is more sensitive to raster degradation than depth. As underlying bathymetric resolution decreases, slope variability reduces. Steep slope areas (>10°) are entirely lost at the coarsest resolution in all regions. Medium slope areas (2–10°) decrease substantially, with reductions of 97.2% in Sabrina Coast, 48.6% in Ross Sea, and 7.5% in the Antarctic Peninsula. In contrast, low-slope areas (0–2°) increase markedly, by 128.5% in SC, 372.5% in RS, and 72.2% in AP (Figure 2, Appendix S1: Figures S2, S3, Table S1). Correlation between the native slope raster and its degraded 4000 m counterpart is moderate to low: Sabrina Coast r = 0.43, Ross Sea r = 0.12, and Antarctic Peninsula r = 0.38, reflecting substantial loss of slope detail (Appendix S1: Figure S5).

### Relationship between Richness and environmental predictor variables at various resolutions

#### Bathymetric terrain attributes as sole predictor variables

Models using only bathymetric terrain attributes (depth, depth², and log-transformed slope) explain a moderate to high amount of variance in species richness for the Sabrina Coast and Ross Sea datasets. For the Antarctic Peninsula dataset, bathymetry alone shows little explanatory power (Figure 3). For the SC dataset, deviance explained (DE) increases as resolution degrades, peaking at 500 m resolution (40.64% DE), before declining to 33.2% DE at 4000 m resolution—an 18.3% decrease in explanatory power. In the Ross Sea dataset, DE increases consistently with coarser resolution, from 26.9% at 40 m to 59.6% at 4000 m—a 121.5% increase. In contrast, all raster resolutions for the Peninsula dataset show low explanatory power using bathymetry alone, with the highest DE at 100 m (9.2%) and the lowest at 4000 m (2.5%), a percentage drop of 72.6%. The ETOPO satellite data performs slightly better than the 4000 m raster for the Peninsula, but slightly worse for the other two regions. A comparative plot of deviance explained values across all resolutions is presented in Figure 3, with detailed percentages in Appendix S1: Table S2.

#### Adding extra environmental predictor variables

When additional environmental predictor variables are included, model performance generally improves. In the Sabrina Coast, DE ranges from 33.5% to 40.6%, which is nearly identical to the terrain-only models. For the Ross Sea, including NPP modestly improves model performance, with DE ranging from 32.0% at 40 m to 59.64% at 4000 m, an 86.3% relative increase. In contrast, the Peninsula dataset experiences a large increase in DE compared to the bathymetry only models, from DE peak at 9.2% (100 m) using only bathymetry to a 52.4% (1000 m) peak when additional environmental variables are included. Model fits remain within a similar range across all resolutions for the Peninsula (46.2%–52.4%). ETOPO-based model predictions perform similarly well to the 4000 m raster in Sabrina Coast and Ross Sea, but marginally better in the Antarctic Peninsula. Although DE fluctuates with changes in bathymetric resolution, the overall differences across resolutions within a region are modest, with the range of DE values remaining relatively narrow. A comparative plot of all deviance explained values (%) for models with added environmental variables is shown in Figure 3 and summarised in Appendix S1: Table S2. A comparative plot of DE values for models fitted individually to each of the three Antarctic Peninsula ecoregions can be found in Appendix S1: Figure S6.

In the Ross Sea, both terrain-only models and those including additional environmental predictors were likely affected by sampling artefacts in the native 40 m bathymetry (Appendix S1: Figure S1), where abrupt depth changes along transect lines produced false steep slopes.

These artefacts smoothed out at coarser resolutions from 2000 m onward. Models potentially influenced by these artefacts are shown in gray in Figure 3.

### Predictive model trends across resolutions

Median predicted richness per cell from models trained and applied at the same resolution ranges from 1.7–1.8 species per cell in the Sabrina Coast (ETOPO: 3.4), 0.9–1.0 in the Ross Sea (ETOPO: 0.8), and 5.4–5.9 in the Antarctic Peninsula (ETOPO: 5.5; Figure 4; Appendix S1: Figures S11, S12).

**Figure 4.**
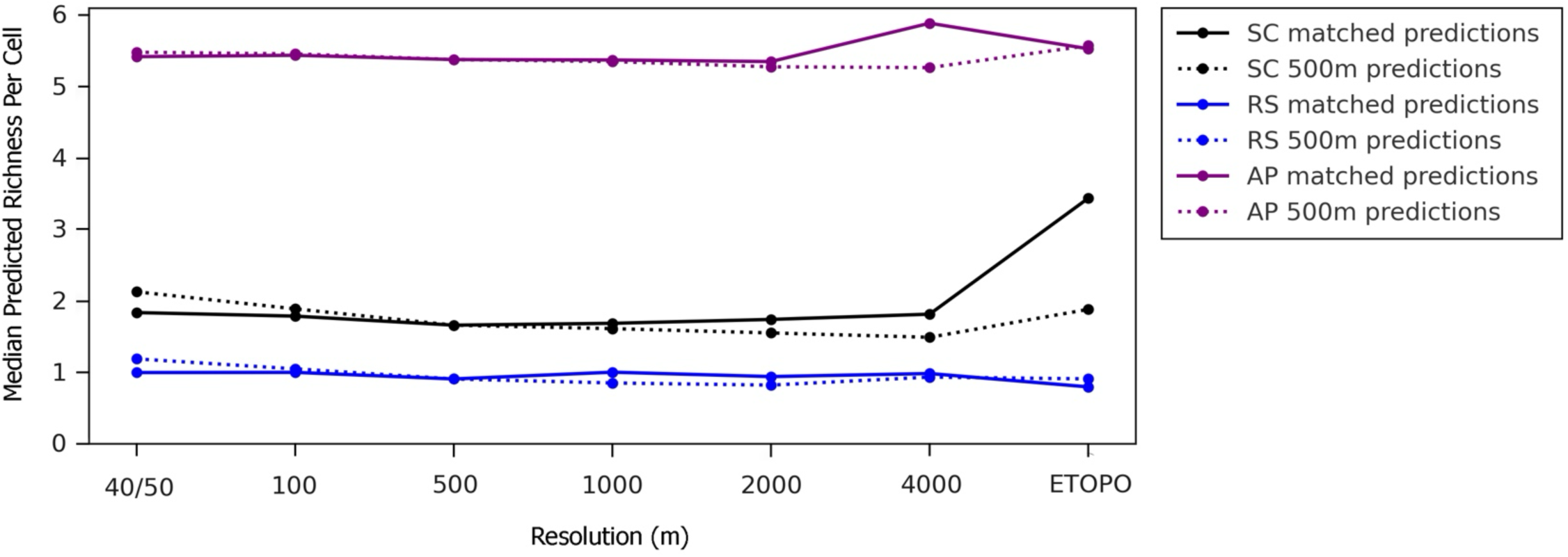
SDM performance when models are applied to resolutions different to the data they were fitted on. Median predicted species richness per cell for models trained and applied at the same resolutions across the Sabrina Coast (SC), Ross Sea (RS), and Antarctic Peninsula (AP), compared to models trained with 500 m data and applied to all other resolutions.

Across regions, models trained on high-resolution data generally underestimate richness when applied to coarser data, whereas models trained on coarser resolutions more often overestimate richness when applied to higher resolution data. When models trained at 40 m resolution are applied to the coarser resolutions, richness is underestimated in 13 of 18 cases and overestimated in 5 of 18 cases (all in the Ross Sea). Underprediction ranges from 0.7% (Antarctic Peninsula 100 m) to 16% (Sabrina Coast 4000 m). In ETOPO-based models, richness is underestimated by 50% in the Sabrina Coast but overestimated by 70% in the Ross Sea. For models trained at 500 m, predictions applied to higher resolutions (40/50 m and 100 m) consistently overestimate richness (6 out of 6 cases), with values ranging from 0.4% (Peninsula 100 m) to 19% (Ross Sea 40 m). When applied to resolutions coarser than 500 m, richness is overestimated in only 2 of 12 cases (both ETOPO), while the remaining 10 models underestimate richness by 0.4–18%, reaching up to 45% underestimation in the Sabrina Coast ETOPO model. In many cases these percentages reflect very small absolute differences in richness values (e.g. the 18% underestimation of the model trained on 500 m bathymetric data and applied to 4000 m data is an absolute richness decrease of 0.32). Trends and absolute mean richness per cell are shown in Figure 4 and Appendix S1: Figures S11, S12.

Species richness prediction maps of the Sabrina Coast region (Figure 5) show that spatial patterns of richness hotspots remain consistent across models, whether trained and applied at 40 m resolution, trained at 40 m and applied to 4000 m data, or trained and applied at 4000 m resolution. While the overall hotspot locations are consistent, predicted richness values per cell differ. In particular, applying the 40 m model to coarser environmental data underestimates richness, with the largest loss occurring in the 99th percentile (Figure 5). Repeating the analyses with percent cover as an alternative biodiversity proxy produced consistent patterns (Appendix S1: Figures S8–S10), reinforcing the robustness of our richness-based results.

**Figure 5.**
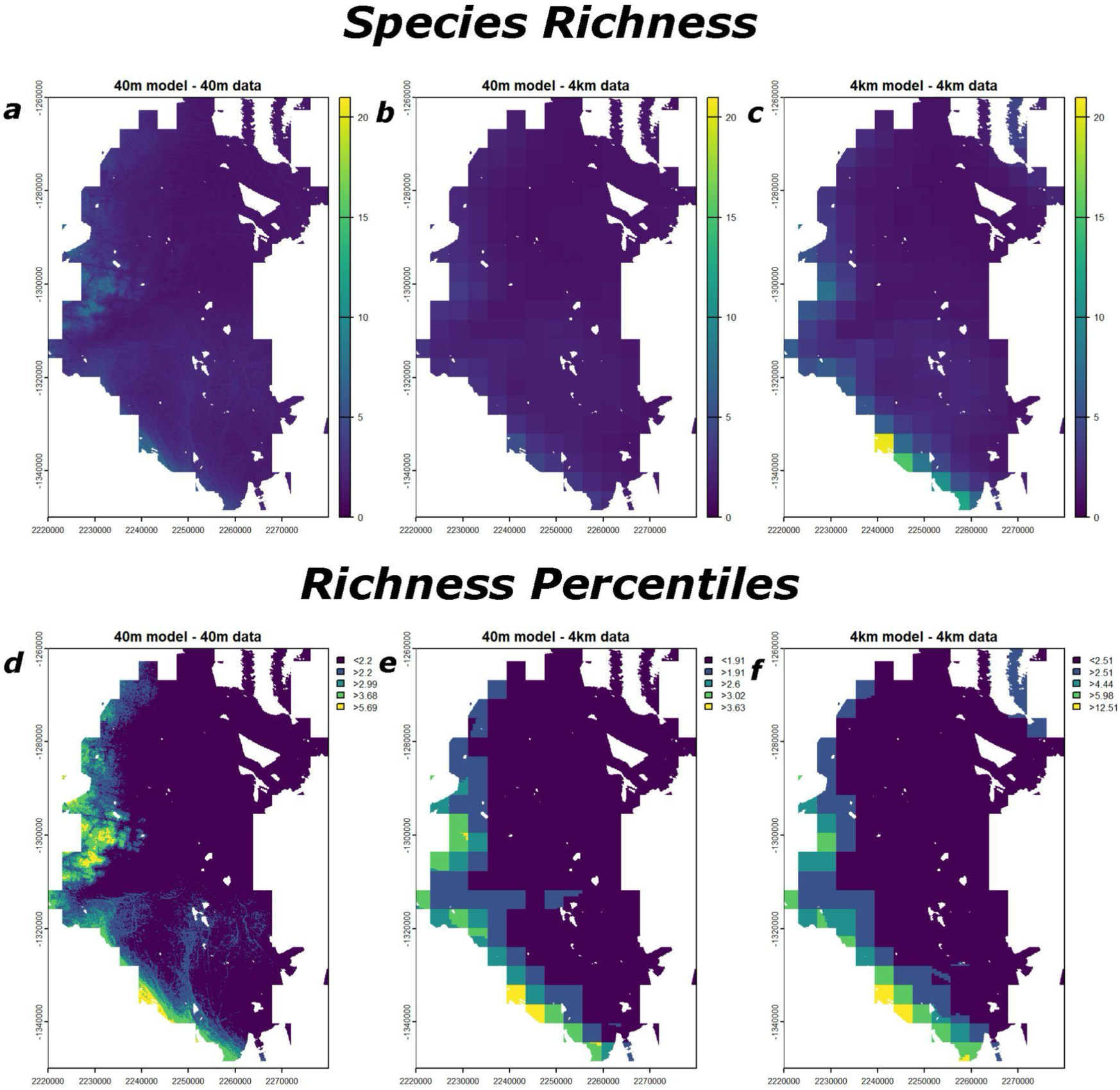
Impact of changing resolutions on richness hotspot locations. Predicted species richness **(a, b, c)** and richness percentiles (75^th^, 90^th^, 95^th^, 99^th^: **d, e, f**) for the Sabrina Coast at 40 m resolution, 4000 m resolution, and the 40 m model applied to 4000 m environmental data. While the spatial distribution of richness hotspots remains consistent across the resolutions, the predicted values change (**d, e, f**), most notably in the 99^th^ percentile.

## Discussion

Our study provides the first quantitative test of how mismatched bathymetric resolutions bias predictions of Antarctic species richness. However, issues can be minimised in practice by matching training and prediction data. When models were trained and applied at the same resolution, predictions of mean richness per cell were highly consistent across the resolution gradient (e.g. SC range: 1.7-1.8). However, mismatched resolutions produced systematic biases. Models trained on 40 m bathymetry tended to underestimate richness (13 out of 18 cases) when applied to coarser resolution data. Models trained at 500 m resolution showed a similar pattern of an underestimation of richness when applied to coarser resolutions (10 out of 12 cases) and overestimated richness (6 of 6 cases) when applied to higher-resolution data. This underestimation was most pronounced in the upper percentiles (95th, 99th), raising concerns about applying models trained on high-resolution bathymetry to unsurveyed areas mapped only at coarse scales. The most likely explanation for this response is that steep slope habitats, which are often associated with high richness (Almond et al. 2021; Post et al. 2022; Smith et al. 2015; De Leo et al. 2014), disappear in coarser-resolution data, leading the model to assume there is less suitable habitat. Similarly, models fitted to coarser-resolution data but predicted into higher-resolution areas will overestimate richness because they encounter more steep-slope habitat than in the training data and may be extrapolating outside the range of the training data. ETOPO-based predictions tended to deviate most strongly, substantially underestimating richness in some regions (e.g. up to 50% in the Sabrina Coast) and overestimating it in others (e.g. up to 70% in the Ross Sea). Given that over 76% of the Southern Ocean is observed solely by satellite-derived bathymetry (Dorschel et al. 2022; Tozer et al. 2019), we expect a high degree of uncertainty associated with predicted richness values in these areas.

Although absolute richness values are sensitive to resolution mismatches between model training and prediction, our results show that spatial patterns of richness hotspots remain remarkably consistent across all resolutions, even when steep-slope detail is lost at coarse scales. This consistency provides confidence that hotspot locations can be reliably identified using coarse-resolution bathymetry when the aim focuses on spatial patterns rather than precise richness counts. However, caution is needed when applying models trained on high-resolution data to coarse-resolution layers from other regions, as this mismatch can lead to systematic under- or overestimation and misinform management decisions. These findings align with broader work in predictive ecology (Meynard et al. 2023; Suárez-Seoane et al. 2014) that underscore the importance of matching bathymetric resolution to study objectives. Biodiversity information is urgently needed to guide spatial planning and policy decisions across the Antarctic continental shelf, but it will likely be many years, if ever, before high-resolution bathymetry is available across the entire region. The minimal variation in deviance explained (DE) across models indicates that model performance remained largely stable, even when trained on coarser bathymetric data. This DE trend along with consistent richness hotspot patterns across resolutions suggests that coarse-scale bathymetry can be robust enough for shelf-wide assessments where broad pattern understanding is more important than absolute richness precision. In practice, there may be a trade-off between increasing confidence in our predictions and generating predictions at a level of enough detail to be practical for management decisions.

As expected, degrading high-resolution bathymetric data to coarser resolutions resulted in a systematic loss of terrain detail. Depth values narrowed in range but remained highly correlated (r > 0.94) between native high-resolution (40/50 m) and degraded 4000 m rasters. In contrast, slope was highly sensitive to degradation: steep slopes (>10°) were entirely lost at 4000 m, and medium slopes (2–10°) were reduced by up to 97% in the Sabrina Coast region, while flat areas (0–2°) expanded by as much as 372% in the Ross Sea. These patterns are consistent with prior work (Post et al. 2020; Dolan and Lucieer 2014) and highlight the vulnerability of slope-dependent terrain metrics to resolution degradation. In this study, where depth remained similar but slope information was substantially lost at coarse resolutions, changes seen in predictive model performance were primarily driven by the loss of slope complexity.

Despite the pronounced smoothing of slope detail, predictive model strength remained more consistent than expected. In the Sabrina Coast, model fit changed little with resolution, regardless of whether only terrain variables or additional predictors (NPP) were included. This suggests that biodiversity patterns are strongly driven by terrain variables in this region and can still be captured using coarse-resolution bathymetry. The Antarctic Peninsula showed a different response: bathymetry alone explained little of the variation in species richness, but model performance improved substantially once the additional predictors of NPP, temperature, and currents were included. This pattern is consistent with earlier work showing that large-scale environmental drivers, such as sea-ice dynamics (and thus NPP) and hydrodynamic characteristics, often outweigh fine-scale terrain features in shaping Antarctic Peninsula communities (Gutt et al. 2016; Henley et al. 2019; Lin et al. 2021). This reinforces the importance of including non-collinear environmental variables to strengthen model reliability, particularly in regions where bathymetry alone has limited explanatory power (Lecours et al. 2015; Peterson and Herkül 2019; Jansen et al. 2019; Pearman et al. 2020). In the Ross Sea, increasing DE with coarser resolution may reflect the influence of larger-scale ecological drivers, although artefacts (Appendix S1: Figure S1) in the original bathymetry likely contributed to lower performance in this case. Together, these findings highlight that adding relevant environmental predictors can buffer against resolution-driven information loss and improve confidence before proceeding with predictive mapping.

While this study offers valuable insight into the role of bathymetric resolution in predictive modelling, it also has limitations. By focusing on overall taxon richness and percent cover, we did not capture potential species interactions and may have masked fine-scale habitat forming species that determine microhabitat assemblages (Katz et al. 2025). Furthermore, observed trends may be region-specific, limiting generalisability across other Antarctic or polar shelves. Future work should align biological sampling with bathymetric scales, incorporate taxon- and trait-specific analyses, and expand to geographically broader regions.

### Conclusion

This study addresses key knowledge gaps in how bathymetric resolution influences predictive habitat suitability models of Antarctic seafloor biodiversity and reveals the biases that arise when models trained on high-resolution bathymetry data are applied to regions mapped only at coarse resolution: richness hotspots hold but counts shift. Both coarse and high-resolution models produce consistent spatial patterns of biodiversity hotspots and similar median richness per cell as long as they are trained and applied at the same resolution. However, applying models trained on high-resolution data to coarse-resolution data led to underestimation of species richness in the majority of cases, particularly in the upper percentiles, reducing confidence in absolute richness values even when hotspot locations remained robust.

Our results underscore the importance of training models on data that match the resolution of the prediction layer to avoid bias. This provides a practical framework for balancing model precision with the realities of patchy data availability, especially in remote, under-sampled regions like the Southern Ocean where >76% of the seafloor is mapped using coarse satellite-derived bathymetry. While high-resolution bathymetric data can be reserved for fine-scale, site-specific management where absolute richness counts are most important, coarse-resolution data may be robust enough for shelf-wide assessment of biodiversity hotspot trends. Continued refinement of predictive methods will be essential for ensuring that conservation policy is guided by models that are both ecologically meaningful and practically scalable.

## Acknowledgements

We thank the many teams who collected, processed, annotated, and made these data publicly available. Sources include the International Bathymetric Chart of the Southern Ocean (IBCSO); NBP1402 and NBP0701 (R/V *Nathaniel B. Palmer*, NCEI); PS81 (R/V *Polarstern*, AWI); and TAN1901 and TAN0802 (R/V *Tangaroa*, NIWA; TAN0802, IPY Census of Antarctic Marine Life).

## Author Contributions

Concept and design of study: SP, JJ, NH, VL

Data analysis: SP, JJ, NH

Visualisation: SP, JJ

Writing - original draft: SP

Writing - review and editing: SP, JJ, NH, VL

## Appendix S1

**Figure S1:**
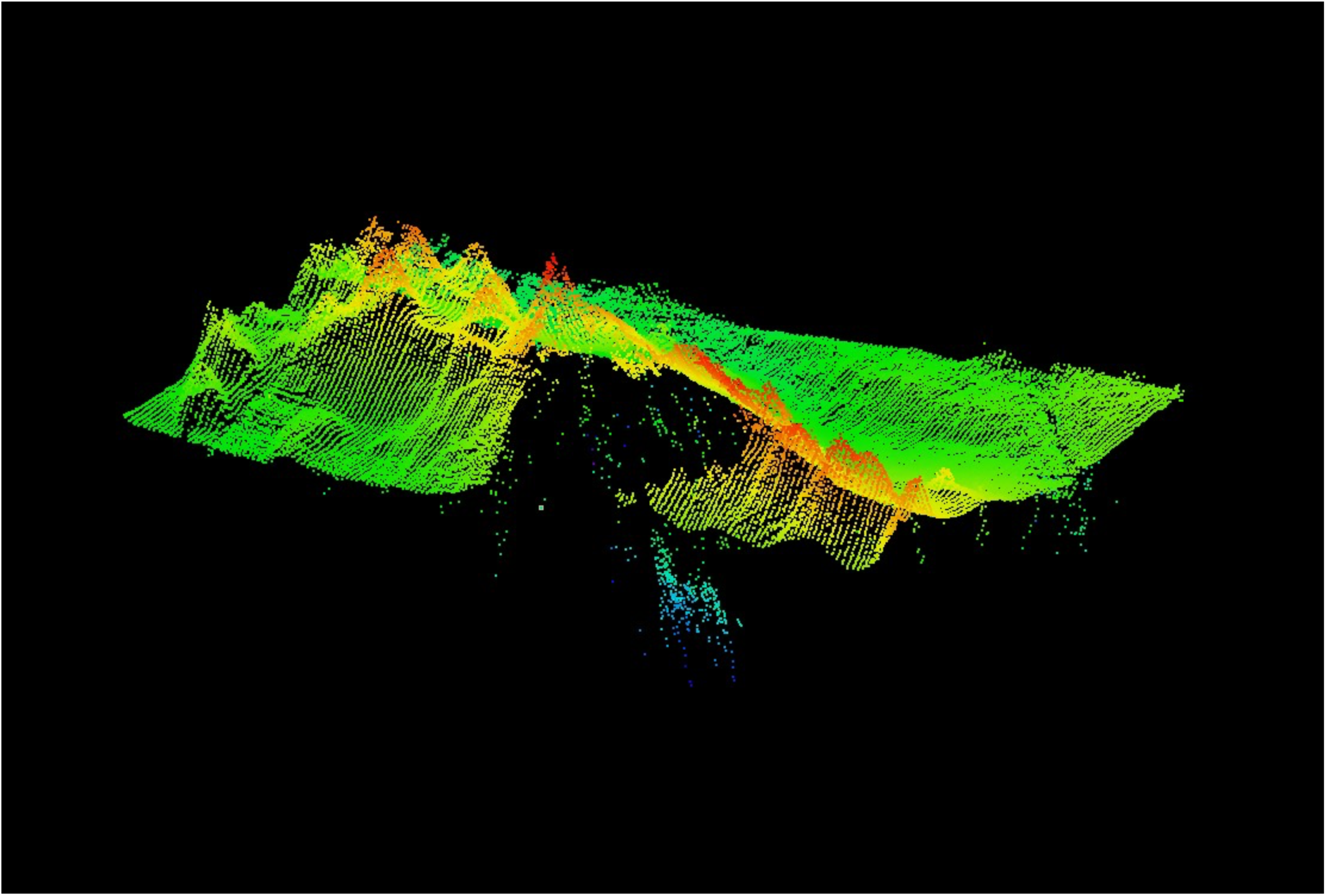
Sampling artefact in Ross Sea 40 m native resolution bathymetry data. The original bathymetric data contained sampling artefacts where abrupt depth changes along some transect lines produced false readings of steep slopes. Note “noise” below sampling plane.

**Figure S2.**
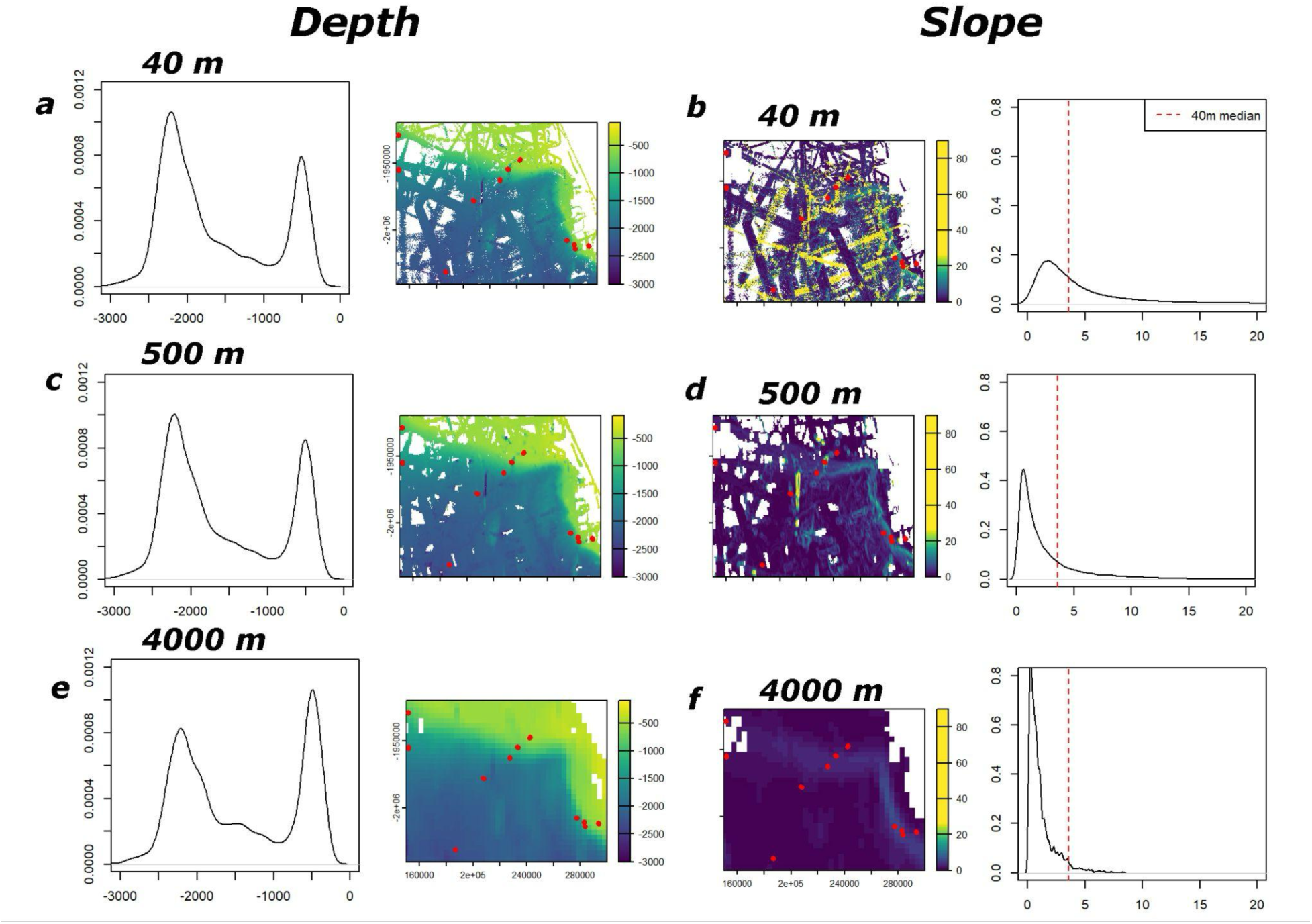
Impact of degrading resolution on depth and slope in the Ross Sea. Terrain variables depth (**left and centre-left column**) and slope (**centre-right and right column**) of the Ross Sea, with density curves (**left and right columns**) comparing the native 40 m raster to aggregated 500 m and 4000 m rasters. Red dots indicate biological sampling locations. Red line on slope density plots (**right column**) represents the median slope value of native 40 m raster. Depth values change minimally, while slope values flatten markedly, especially at slopes >10°.

**Figure S3.**
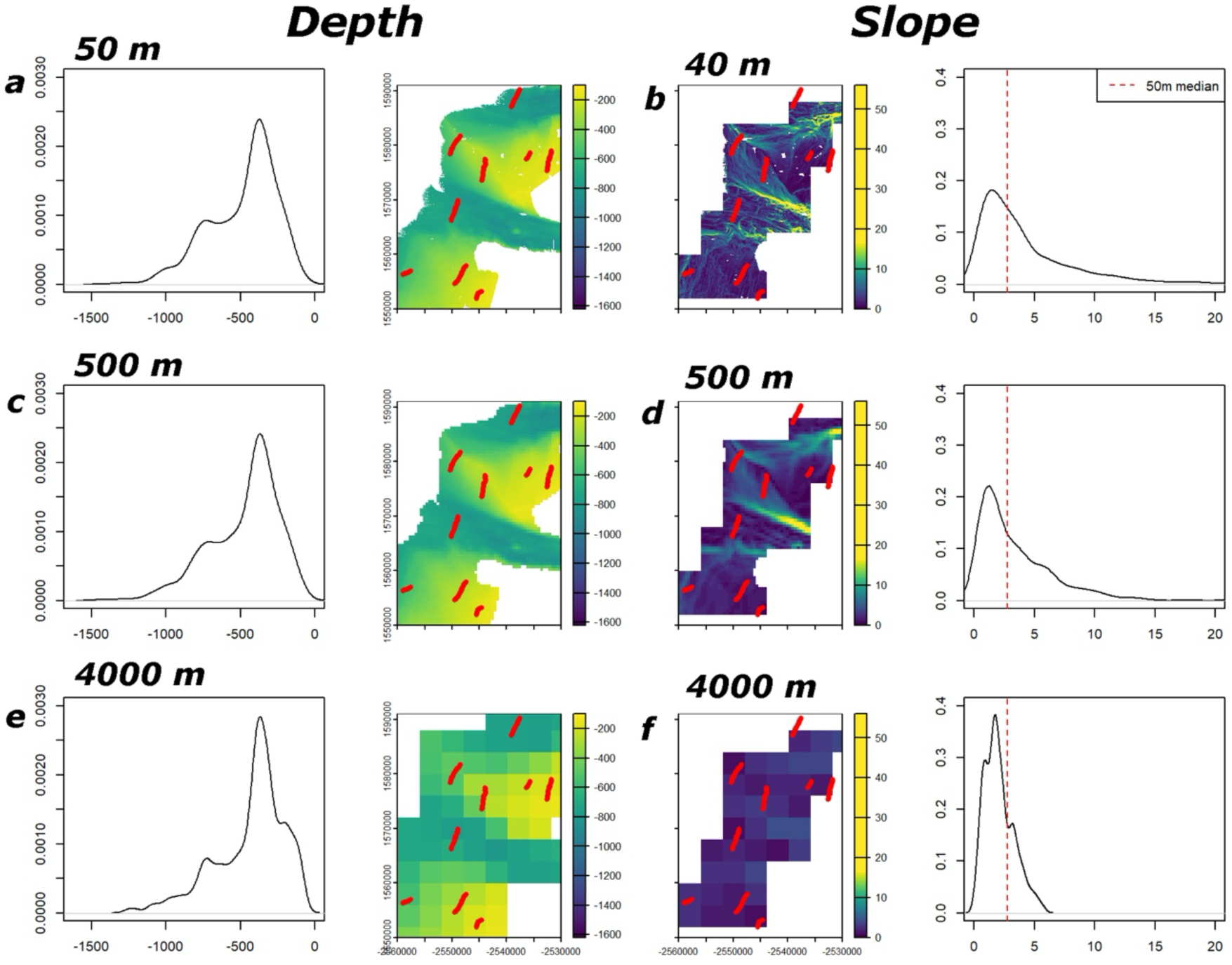
Impact of degrading resolution on depth and slope of the Antarctic Peninsula. Terrain variables depth (**left and centre-left column**) and slope (**centre-right and right column**) of the Antarctic Peninsula, with density curves (**left and right columns**) comparing the native 50 m raster to aggregated 500 m and 4000 m rasters. Red dots indicate biological sampling locations. Red line on slope density plots (**right column**) represents the median slope value of native 50 m raster. Depth values change minimally, while slope values flatten markedly, especially at slopes >10°. Due to the large extent of the peninsula dataset, just a small portion of the Bransfield Strait section is plotted on the map for visual clarity, however density plots reflect the entire Peninsula extent.

**Figure S4.**
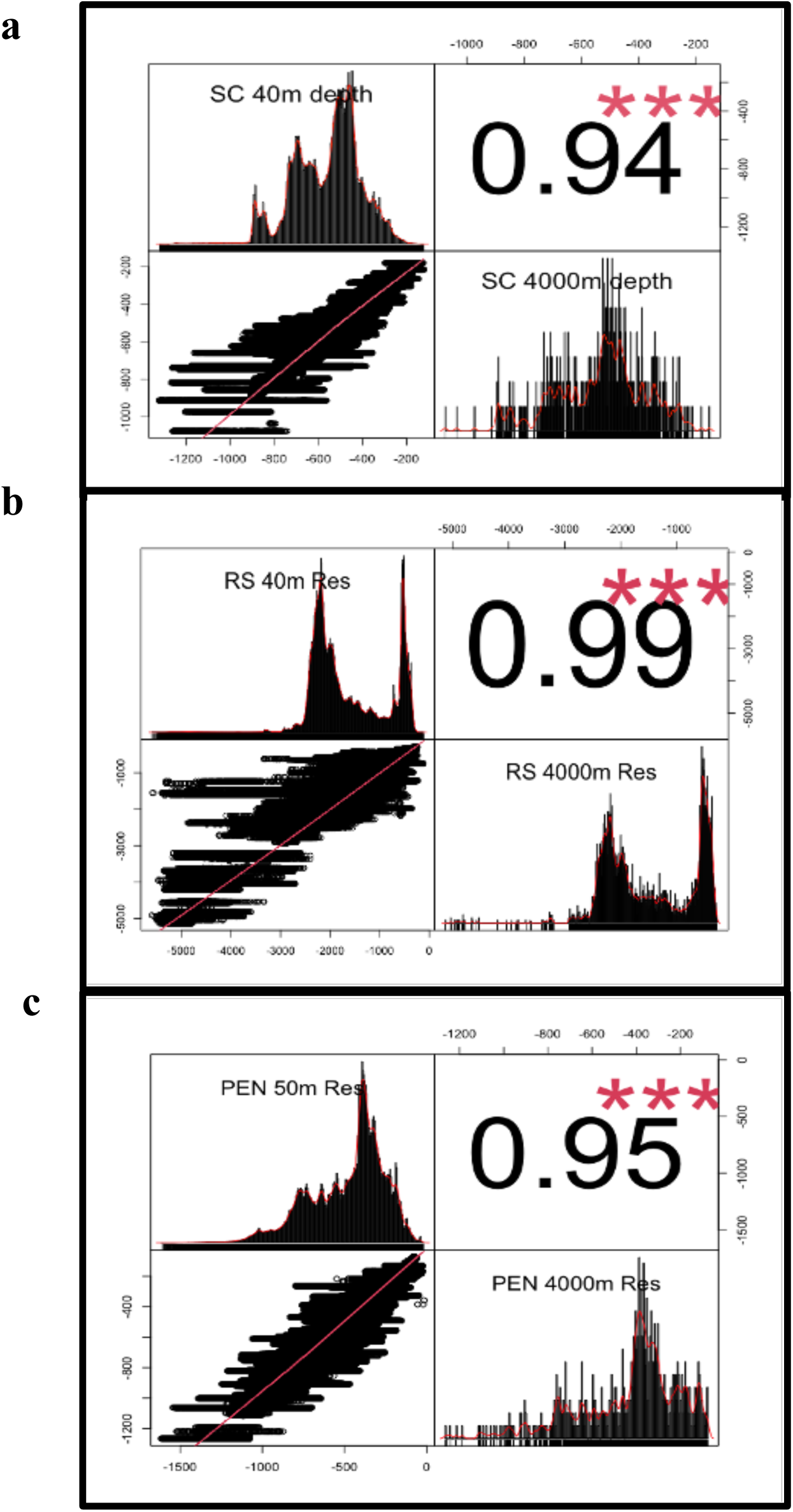
Depth Biplots. Sabrina Coast (**a**), Ross Sea (**b**) and Antarctic Peninsula (**c**) native 40/50 m depth rasters and the degraded 4000 m depth rasters with associated histograms, absolute values of Pearson’s correlation between variables and bivariate scatter plot with fitted line.

**Figure S5.**
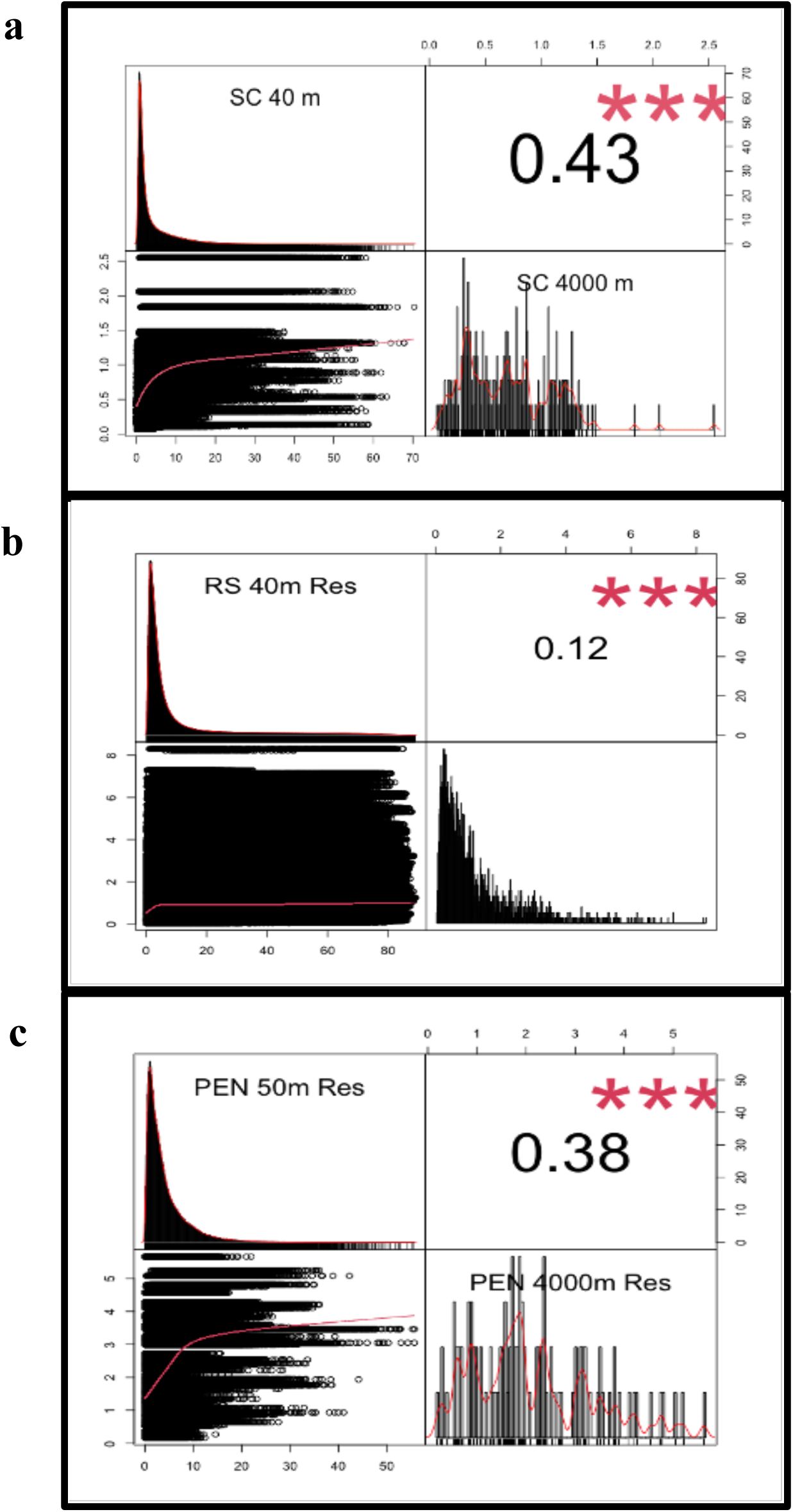
Slope Biplots. Sabrina Coast (**a**), Ross Sea (**b**) and Antarctic Peninsula (**c**) native 40/50 m slope rasters and the degraded 4000 m slope rasters with histogram, absolute values of Pearson’s correlation between variables and bivariate scatter plot with fitted line.

**Figure S6.**
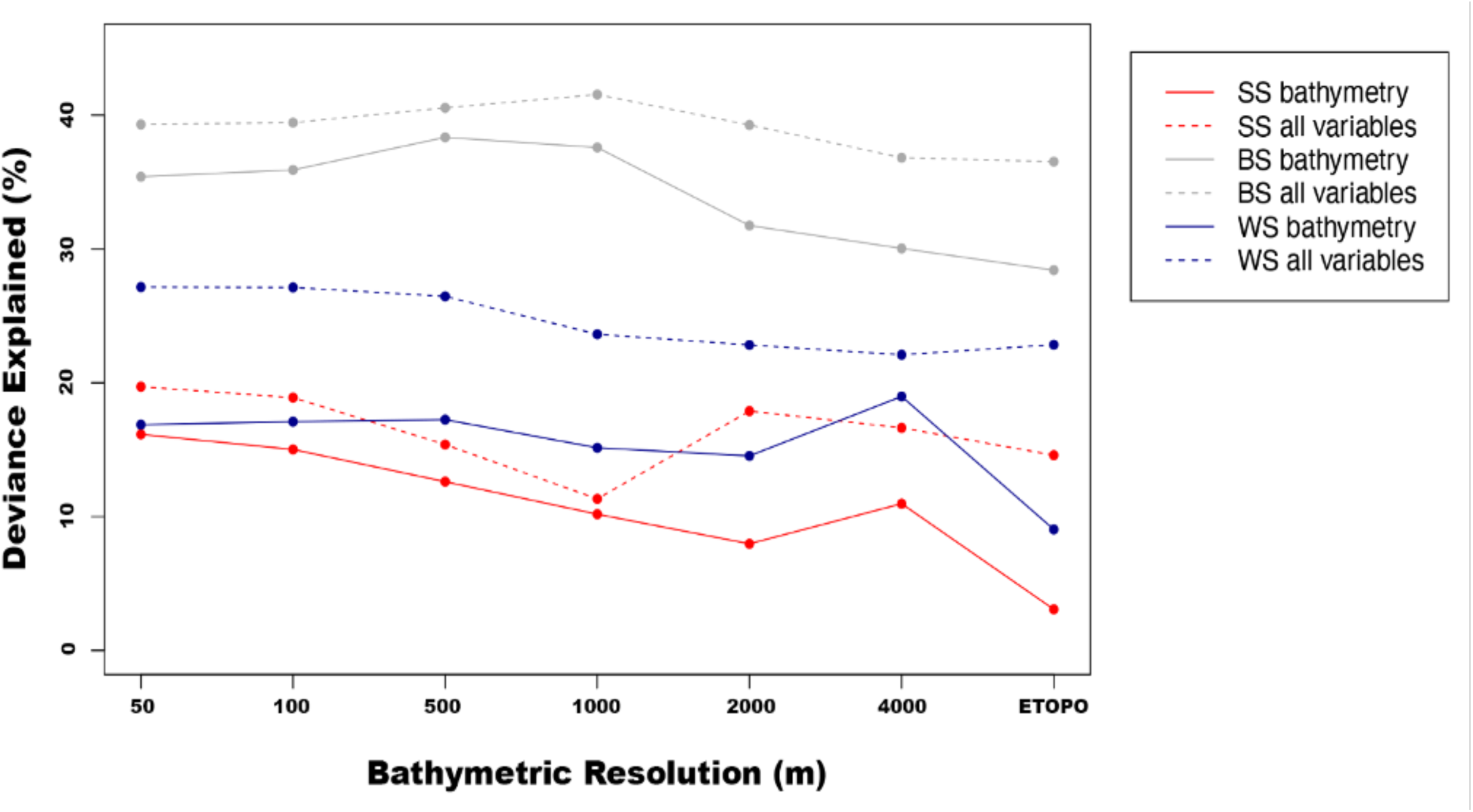
Deviance explained (DE, %) across raster resolutions for Antarctic Peninsula ecoregions South Shetland (SS), Bransfield Strait (BS), and Weddell Sea (WS) from generalized linear models. Results are shown for bathymetry-only predictors and for models including additional environmental predictors. DE drops markedly for the ETOPO raster in SS and WS under bathymetry-only models, but this decline largely disappears when environmental predictors are added. Model strength for all regions generally increases with the added environmental variables.

**Figure S7.**
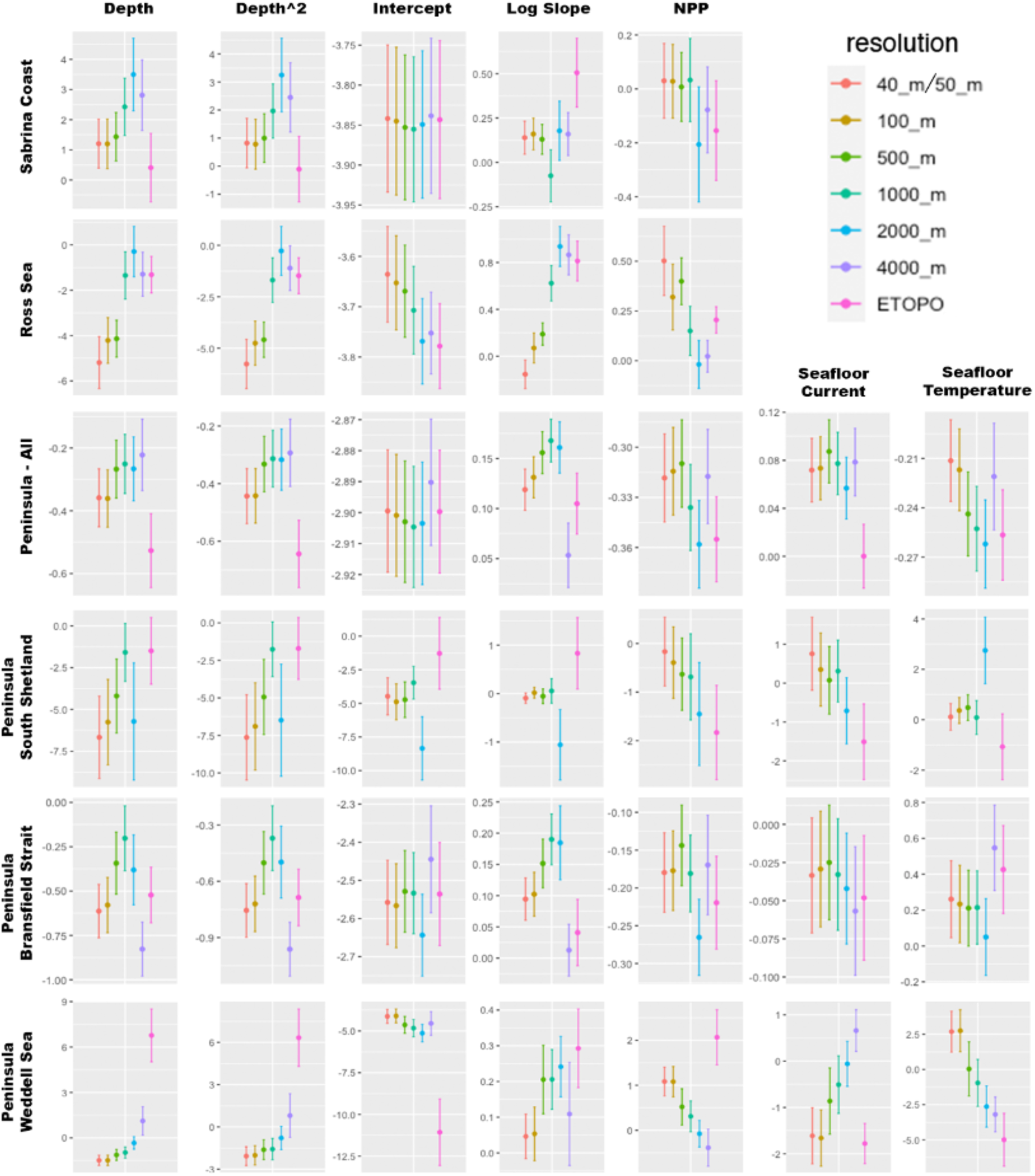
Coefficient estimate outputs (+/- SE) from each generalized linear model. Log-scale models with all variables included plotted for all study regions at each raster resolution. The plotted coefficients allow visualisation of the relationship between each predictor variable and species richness, and how that relationship changes as the bathymetric resolution decreases. In the Peninsula South Shetland dataset, none of the variables from the 4000 m raster show a relationship with richness, which is why they are excluded from the plot.

**Figure S8.**
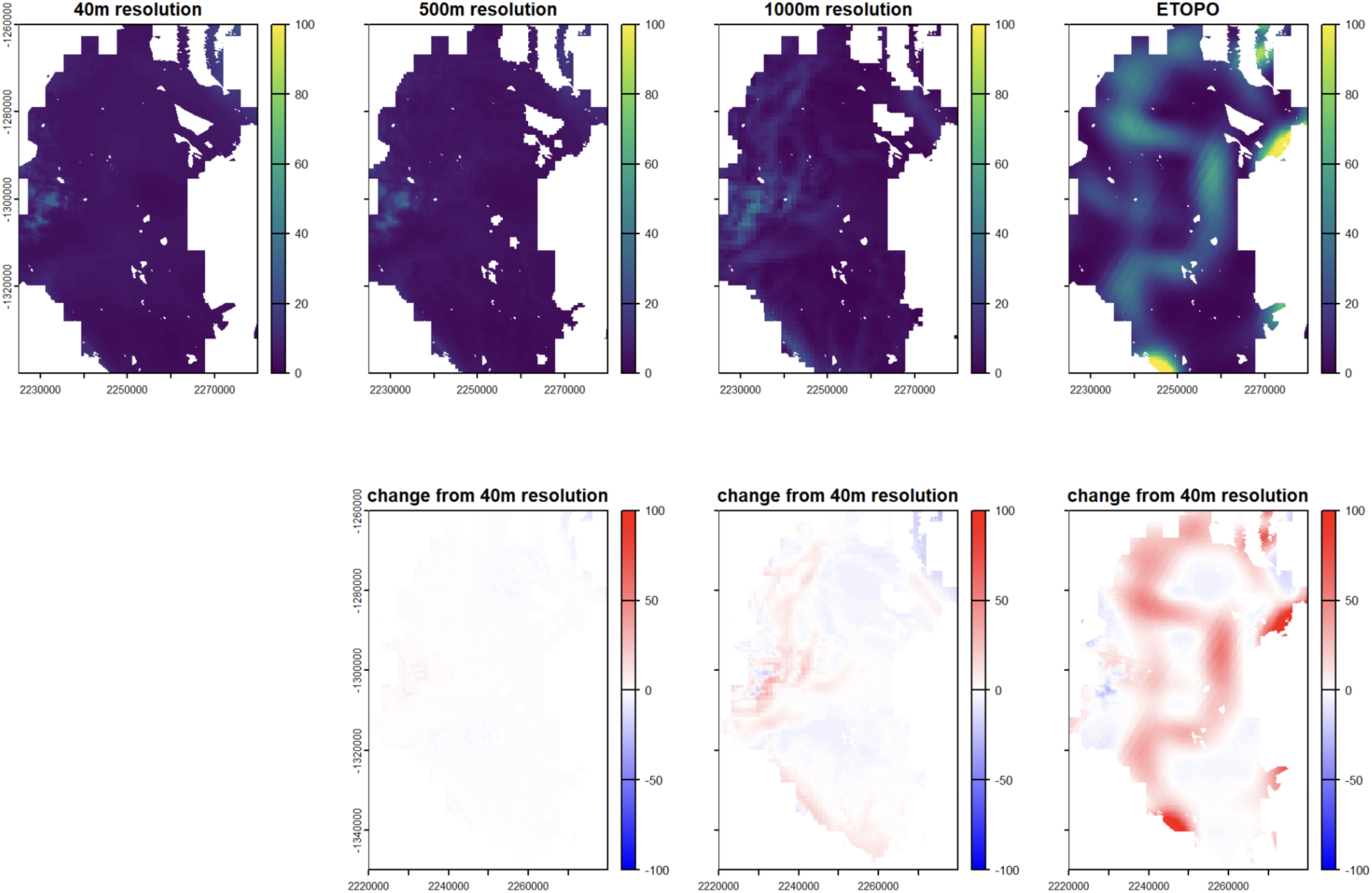
Predicted Percent Cover for the Sabrina Coast. 40 m, 500 m, 1000 m, and ETOPO resolutions with difference plots compared to the 40 m raster. Note that the spatial distribution of percent cover hotspots remains consistent across the resolutions.

**Figure S9.**
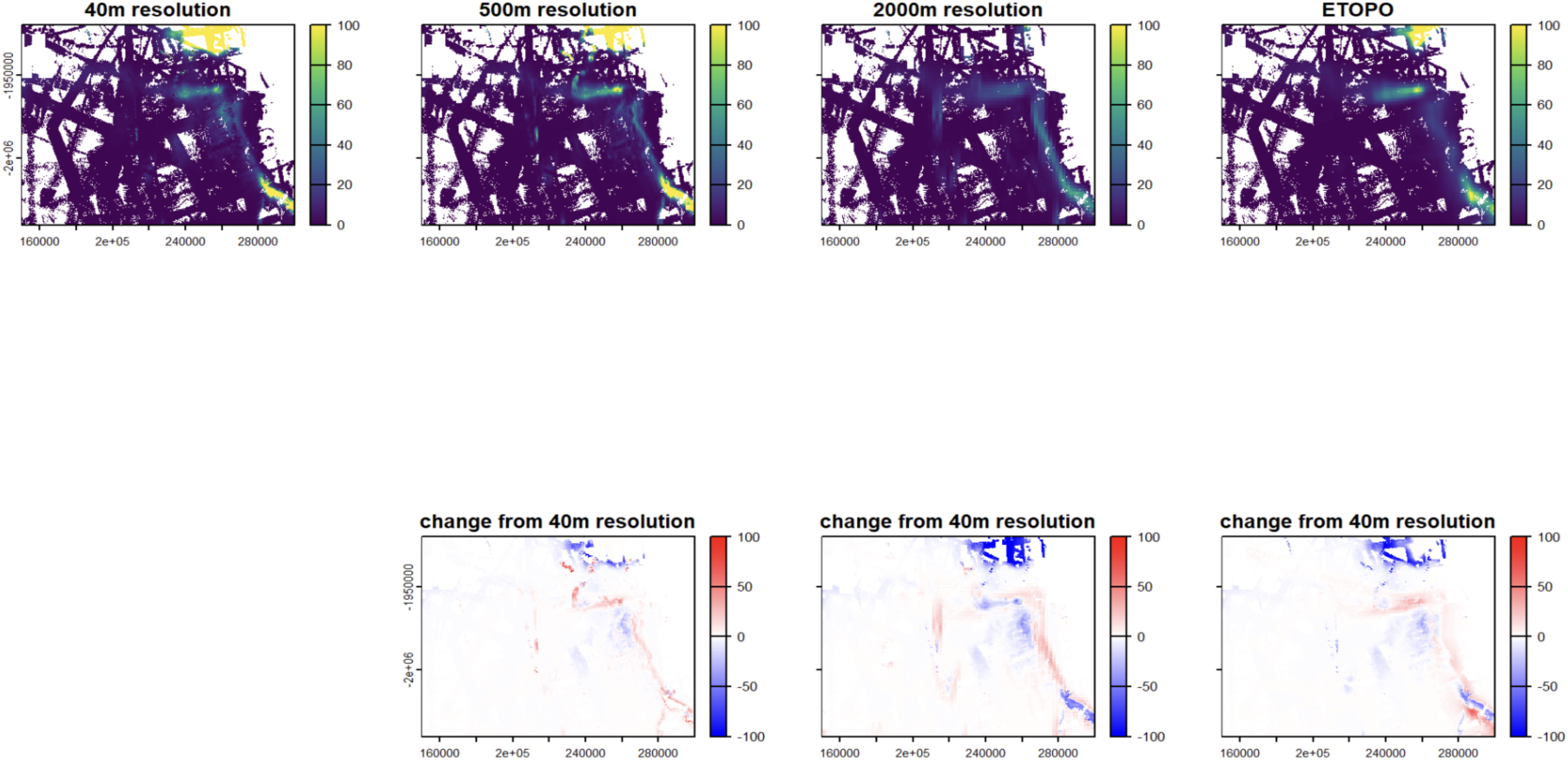
Predicted Percent Cover for the Ross Sea. 40 m, 500 m, 1000 m, and ETOPO resolutions with difference plots compared to the 40 m raster. Note that the spatial distribution of percent cover hotspots remains consistent across the resolutions.

**Figure S10.**
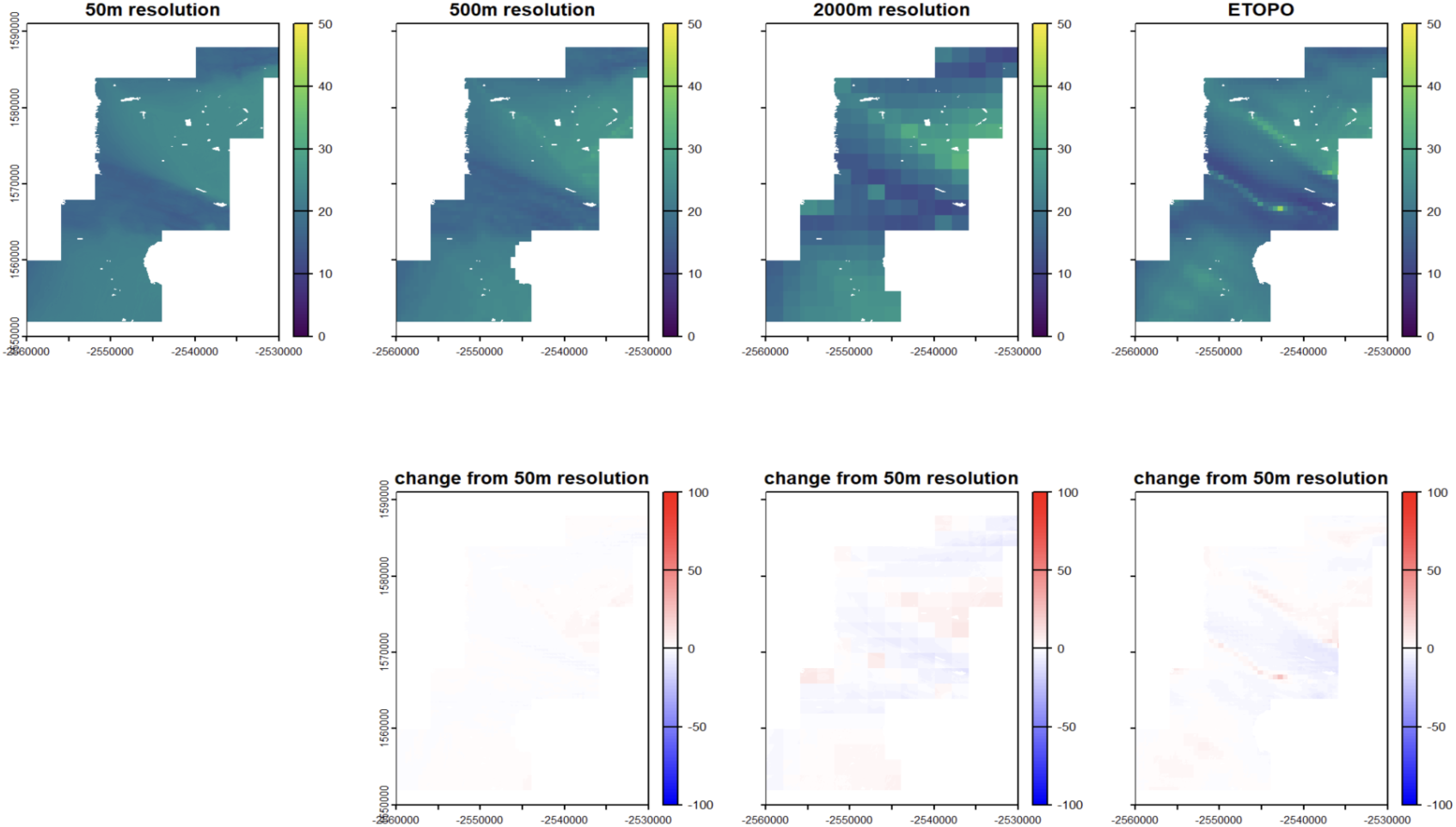
Predicted Percent Cover for the Antarctic Peninsula. 40 m, 500 m, 1000 m, and ETOPO resolutions with difference plots compared to the 40 m raster. Note that the spatial distribution of percent cover hotspots remains consistent across the resolutions.

**Figure S11.**
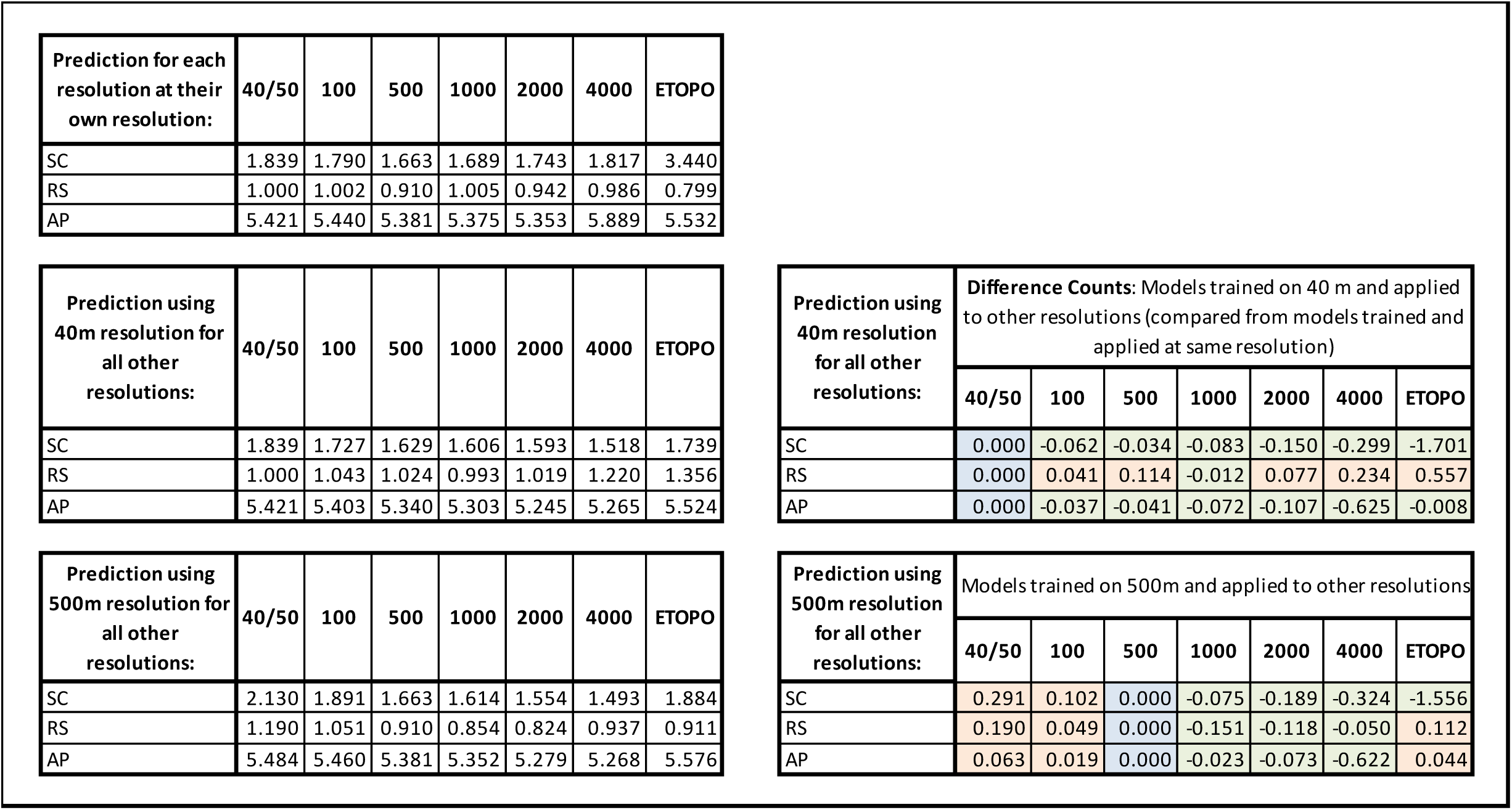
Mean richness predictions across resolutions and regions. Mean predicted richness per cell for models trained and applied at their own resolutions, and for models trained at 40/50 m and 500 m then applied to all other resolutions. Coloured tables show prediction differences in absolute richness counts for the 40/50 m trained models and 500 m trained models compared to the matching resolution models. Blue shows models trained and predicted at matching resolutions, orange shows overestimations of richness, green shows underestimation of richness.

**Figure S12.**
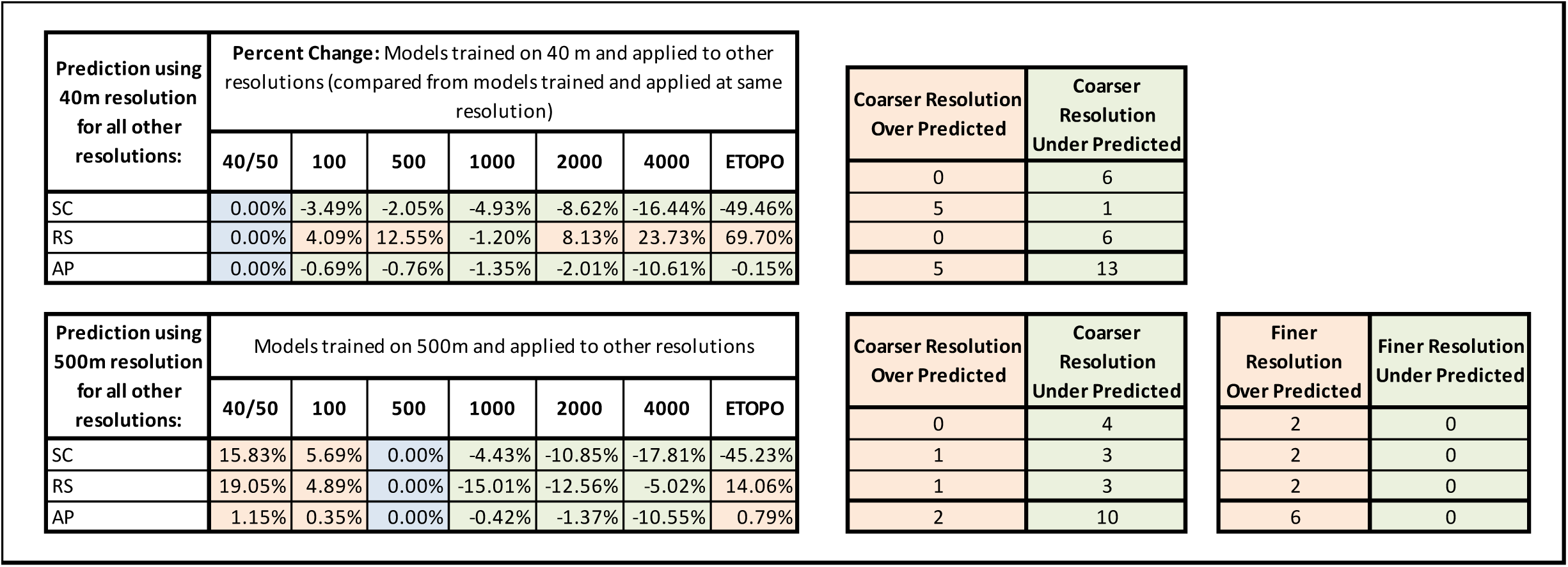
Richness percent change across resolutions and regions. Richness percent change of models trained at 40/50 m resolution applied to all other resolutions, and models trained on 500 m applied to all other resolutions (compared to models trained and applied at matching resolutions). Counts of over- and underestimation per region shown on right. Blue shows models trained and predicted at matching resolutions, orange shows overestimations of richness, green shows underestimation of richness.

**Table S1.** A comprehensive overview of calculated variables for all original bathymetric rasters and their aggregated coarser rasters. Variables include: Depth max (m), depth min (m), depth mean (m), slope min (degrees), slope max (degrees), slope mean (degrees), observations in slope category 0-2 degrees, observations in slope category 2-10 degrees, observations in slope category 10-100 degrees, percent change of slope in category 0-2 degrees, percent change in slope category 2-10 degrees, and percent change in slope category 10-100 degrees.

| Rasters & Resolutions | Depth Max (m) | Depth Min (m) | Depth Mean (m) | Slope Min (deg) | Slope Max (deg) | Slope Mean (deg) | Slope value number 0-2 ° | Slope value numbers 2-10 ° | Slope value numbers 10-100 ° | % chng 0-2 ° | % chng 2-10 ° | % chng 10-100 ° |
| --- | --- | --- | --- | --- | --- | --- | --- | --- | --- | --- | --- | --- |
| SC 40m | 1321 | 121 | 560 | 0 | 70 | 4 | 983624 | 674141 | 205269 | 0 | 0 | 0 |
| SC 100m | 1318 | 127 | 560 | 0 | 44 | 3 | 1154737 | 628408 | 74006 | 17.4 | -6.8 | -63.9 |
| SC 500m | 1245 | 143 | 556 | 0 | 24 | 1 | 1434900 | 376836 | 14172 | 45.9 | -44 | -93 |
| SC 1000m | 1222 | 159 | 552 | 0 | 14 | 1 | 1598751 | 208076 | 8471 | 62.5 | -69.1 | -95.9 |
| SC 2000m | 1076 | 161 | 545 | 0 | 5 | 1 | 1720954 | 74994 | 0 | 74.9 | -88.9 | -100 |
| SC 4000m | 1076 | 153 | 533 | 0 | 3 | 1 | 2247336 | 19208 | 0 | 128.47 | -97 | -100 |
| RS 40m | 5597 | 114 | 1622 | 0 | 89 | 10 | 4761296 | 8553985 | 3865122 | 0 | 0 | 0 |
| RS 100m | 5597 | 120 | 1630 | 0 | 85 | 6 | 9032482 | 6546920 | 2977892 | 89.7 | -23.5 | -23 |
| RS 500m | 5458 | 179 | 1594 | 0 | 66 | 3 | 11768710 | 6448690 | 857200 | 147.2 | -24.6 | -77.8 |
| RS 1000m | 5458 | 260 | 1556 | 0 | 46 | 2 | 13575324 | 5462531 | 421428 | 185.1 | -36 | -89.1 |
| RS 2000m | 5163 | 270 | 1509 | 0 | 13 | 2 | 16451516 | 5077650 | 69126 | 245.5 | -40.6 | -98.2 |
| RS 4000m | 5128 | 279 | 1476 | 0 | 8 | 1 | 22496211 | 4393206 | 0 | 372.5 | -48.6 | -100 |
| PEN 50m | 1615 | 16 | 483 | 0 | 56 | 4 | 215585 | 304270 | 50634 | 0 | 0 | 0 |
| PEN 100m | 1606 | 16 | 483 | 0 | 45 | 4 | 229476 | 301540 | 45252 | 6.4 | -0.9 | -10.6 |
| PEN 500m | 1573 | 17 | 473 | 0 | 21 | 3 | 261100 | 290900 | 23800 | 21.1 | -4.4 | -53 |
| PEN 1000m | 1509 | 35 | 462 | 0 | 15 | 3 | 261600 | 300800 | 7600 | 21.3 | -1.1 | -84 |
| PEN 2000m | 1417 | 38 | 449 | 0 | 9 | 3 | 264000 | 310400 | 0 | 22.5 | 2 | -100 |
| PEN 4000m | 1266 | 75 | 433 | 0 | 6 | 2 | 371200 | 281600 | 0 | 72.2 | -7.5 | -100 |

**Table S2.**
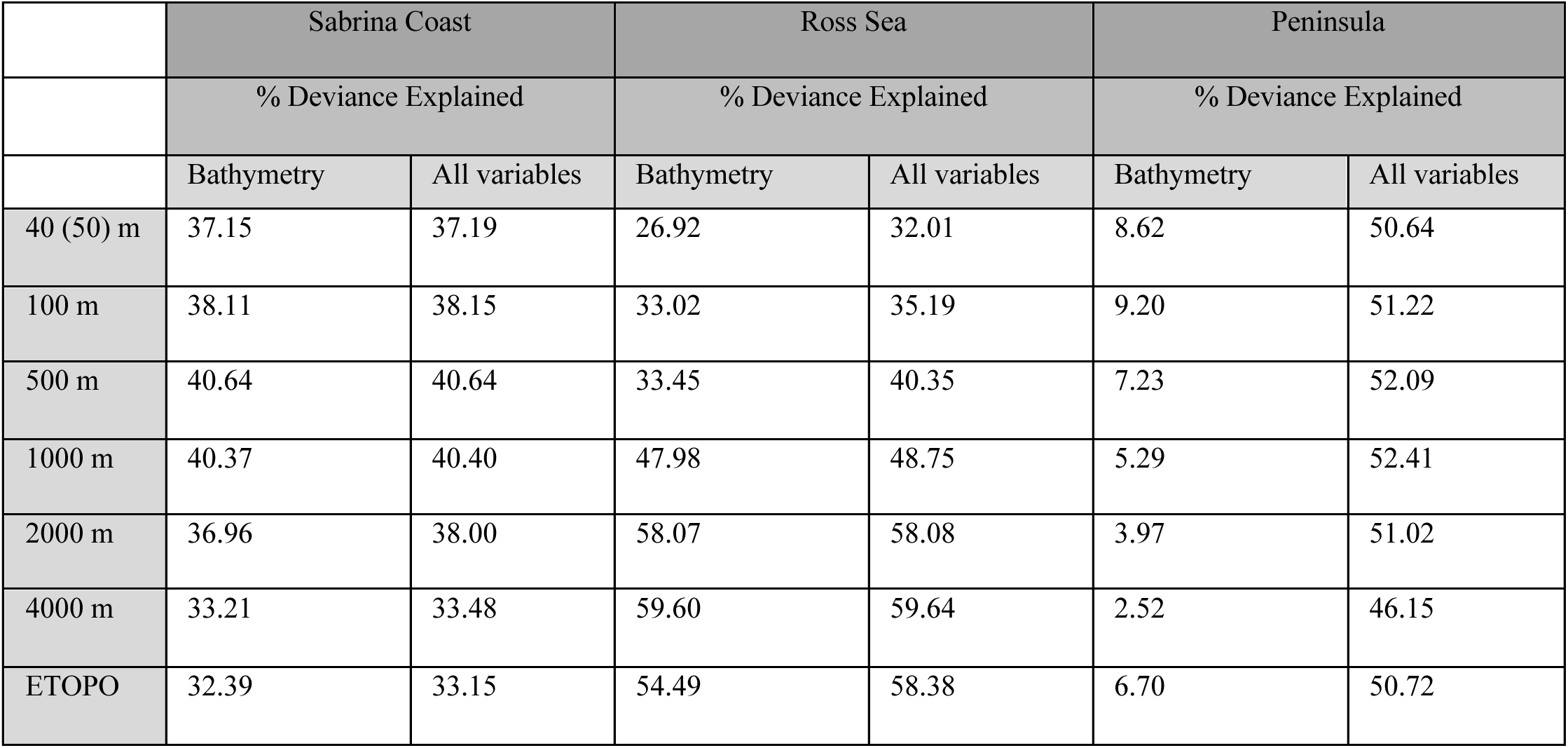
Deviance explained (%). DE for bathymetry-only versus all environmental predictors for all raster resolutions across Sabrina Coast, Ross Sea, and Antarctic Peninsula. Native resolution is 40 m for Sabrina Coast and Ross Sea, 50 m for Antarctic Peninsula.

|  | Sabrina Coast |  | Ross Sea |  | Peninsula |  |
| --- | --- | --- | --- | --- | --- | --- |
|  | % Deviance Explained |  | % Deviance Explained |  | % Deviance Explained |  |
|  | Bathymetry | All variables | Bathymetry | All variables | Bathymetry | All variables |
| 40 (50) m | 37.15 | 37.19 | 26.92 | 32.01 | 8.62 | 50.64 |
| 100 m | 38.11 | 38.15 | 33.02 | 35.19 | 9.20 | 51.22 |
| 500 m | 40.64 | 40.64 | 33.45 | 40.35 | 7.23 | 52.09 |
| 1000 m | 40.37 | 40.40 | 47.98 | 48.75 | 5.29 | 52.41 |
| 2000 m | 36.96 | 38.00 | 58.07 | 58.08 | 3.97 | 51.02 |
| 4000 m | 33.21 | 33.48 | 59.60 | 59.64 | 2.52 | 46.15 |
| ETOPO | 32.39 | 33.15 | 54.49 | 58.38 | 6.70 | 50.72 |

